# Capturing spatiotemporal variation in salt marsh belowground biomass, a key resilience metric, through geoinformatics

**DOI:** 10.1101/2024.09.16.613282

**Authors:** Kyle D Runion, Deepak R Mishra, Merryl Alber, Mark A Lever, Jessica L O’Connell

## Abstract

The Belowground Ecosystem Resiliency Model (BERM) is a geoinformatics tool that was developed to predict belowground biomass (BGB) of *Spartina alterniflora* in salt marshes based on remote sensing of aboveground characteristics and other readily available hydrologic, climatic, and physical data. We sought to characterize variation in *S. alterniflora* BGB over both temporal and spatial gradients through extensive marsh field observations in coastal Georgia, USA, to quantify their relationship with a suite of predictor variables, and to use these results to improve performance and expand the parameter space of BERM. We conducted pairwise comparisons of *S. alterniflora* growth metrics measured at nine sites over three to eight years and found that BGB grouped by site differed in 69% of comparisons, while only in 21% when grouped by year. This suggests that BGB varies more spatially than temporally. We used the BERM machine learning algorithms to evaluate how variables relating to biological, climatic, hydrologic, and physical attributes covaried with these BGB observations. Flooding frequency and intensity were most influential in predicting BGB, with predictor variables related to hydrology composing 61% of the total feature importance in the BERM framework. When we used this expanded calibration dataset and associated predictors to advance BERM, model error was reduced from a normalized root mean square error of 13.0% to 9.4% in comparison to the original BERM formulation. This reflects both an improvement in predictive performance and an expansion in conditions for potential model application. Finally, we used regression commonality analysis to show that model estimates reflected the spatiotemporal structure of BGB variation observed in field measurements. These results can help guide future data collection efforts to describe landscape-scale BGB trends. The advanced BERM is a robust tool that can characterize *S. alterniflora* productivity and resilience over broad spatial and temporal scales.

## 1. Introduction

Salt marshes are vulnerable habitats that are increasingly threatened by sea level rise (SLR) (Morris et al. 2002, Craft et al. 2009, Crosby et al. 2016). Belowground plant production and the accumulation of organic matter builds soil volume and elevation, contributing to vertical accretion, carbon storage, and marsh resilience (Turner et al. 2002, Turner et al. 2004, Gonneea et al. 2019). Belowground biomass (BGB) is therefore an important indicator of salt marsh resilience and susceptibility to global change. However, belowground biomass has high spatiotemporal variability (Gallagher 1983, Mendelssohn and Morris 2000, O’Connell et al. 2020), making it challenging to develop geospatial tools that can accurately predict BGB at the landscape scale. In this paper, we used an extended dataset of observations collected in *Spartina alterniflora*-dominated marshes to characterize spatiotemporal variation in BGB and advance the Belowground Ecosystem Resiliency Model (BERM).

*Spartina alterniflora* is the dominant form of vegetation in many low marsh ecosystems across the U.S. Atlantic and Gulf Coasts (Pennings and Bertness 2001). Although it is extremely well studied (Pyšek et al. 2008), BGB has received far less attention than aboveground biomass (AGB) (Rivera-Monroy et al. 2019). AGB of *S. alterniflora* varies with environmental gradients, with taller plants generally located nearer the creek edge and shorter plants in the marsh interior (Mendelssohn and Morris 2000). These plants have different phenologies along the creek to interior gradient (O’Connell et al. 2020), and are likely to have different belowground characteristics. Studies have found short-form *S. alterniflora* to have greater BGB than tall-form plants, and it has been suggested that low redox potential and high soil sulfides in the marsh interior may drive this difference in plant growth and biomass allocation (Gallagher et al. 1988, Ornes and Kaplan 1989, Blum 1993). However, BGB samples are difficult to collect and labor-intensive to analyze, making it challenging to resolve spatial and temporal patterns at the landscape scale. Spatially, BGB has been found to differ over small biophysical and hydrologic gradients such as elevation, soil temperature, and flooding regime (Snedden et al. 2015, Hanson et al. 2016, Crosby et al. 2017, O’Connell et al. 2021). This can lead to heterogeneity across short distances and among marsh units. Temporally, plant biomass exhibits strong seasonality, with belowground carbohydrate reserves utilized in early spring to stimulate aboveground production and replenished during the growing season to a maximum belowground stock prior to winter (Gallagher 1983, Jung and Burd 2017). Interannual variation may also be substantial (Darby and Turner 2008a), but such data records are sparse (Rivera-Monroy et al. 2019). The relationship between BGB and environmental drivers is also unclear.

The Belowground Ecosystem Resiliency Model (BERM) is a recently developed geospatial informatics tool to characterize salt marsh resilience and productivity in near real-time, initially calibrated in U.S. Georgia (GA) coast *S. alterniflora* marshes (O’Connell et al. 2021). The main goal of BERM is to estimate *S. alterniflora* total BGB. BERM integrates empirical measurements of *S. alterniflora* BGB with remotely sensed aboveground biophysical metrics, climate variables, elevation, and data from tide stations. BERM was built from an extreme gradient boosting model framework and uses the ‘xgboost’ package in R (Chen et al. 2019). Estimates are made at a monthly timestep and 30 m spatial resolution, which allows us to reconstruct spatiotemporal variation in response to past conditions and provide near real-time estimates of BGB derived from satellite and other gridded spatial data. While BERM was a step forward for evaluating *S. alterniflora* BGB, a limited calibration dataset resulted in a narrow parameter space. The calibration dataset of O’Connell et al. (2021) consisted of data collected at four different salt marsh sites. Data at three of these sites were limited to a single growing season, and thus lacked the temporal coverage necessary to portray intra- and interannual variation. Further, the spatial coverage was small; two of four sites were located ∼1.2 km apart within the same marsh unit, effectively resulting in three distinct marsh sites. This paper broadens BERM applicability by expanding the calibration dataset to include ground-truth data that exhibit a wider range of hydrodynamic and biogeochemical settings. In the process, we also sought to understand which environmental gradients best describe BGB variation. Our goal with this is to provide a tool to inform coastal management and salt marsh conservation and restoration.

The overall goal of this study was to improve our understanding of the dynamics of BGB of *S. alterniflora* and how it relates to environmental drivers. We used extensive field measurements in combination with BERM to 1) compare spatial and temporal scales of BGB variability and 2) identify relevant environmental predictors of BGB, 3) recalibrate and update BERM for a broad range of environmental conditions, and 4) evaluate the spatiotemporal structure of model estimates. Our results can be used to identify areas that are potentially vulnerable to SLR, and to guide future data collection to calibrate BERM so that it can be applied in other areas.

## 2. Methods

### 2.1. Study Area

Our study area included *S. alterniflora*-dominated marshes mostly within the GA Coastal Ecosystems – Long Term Ecological Research (GCE-LTER) project domain (∼1,000 km^2^) in coastal GA, USA (Figure 1a, 1b). Study sites were concentrated near Sapelo Island, GA, where diurnal tides have a mean tidal range of ∼2 m, and mean sea level (MSL) is −0.07 m NAVD88 (at the nearby Fort Pulaski NOAA CO-OPS station). On Sapelo Island, marshes have remained in mostly pristine condition due to relatively low nutrient loading from the watershed (Bricker et al. 2007, Schaefer and Alber 2007, Weston et al. 2009) and restricted public access to the island (Sapelo Island National Estuarine Research Reserve 2008). Urban development on Sapelo Island is minimal; a small residential community, Hog Hammock, along with the tourism industry and research activities, result in a population density of ∼5 people per square mile (Sullivan 2019, McLachlan et al. 2021).

**Figure 1.**
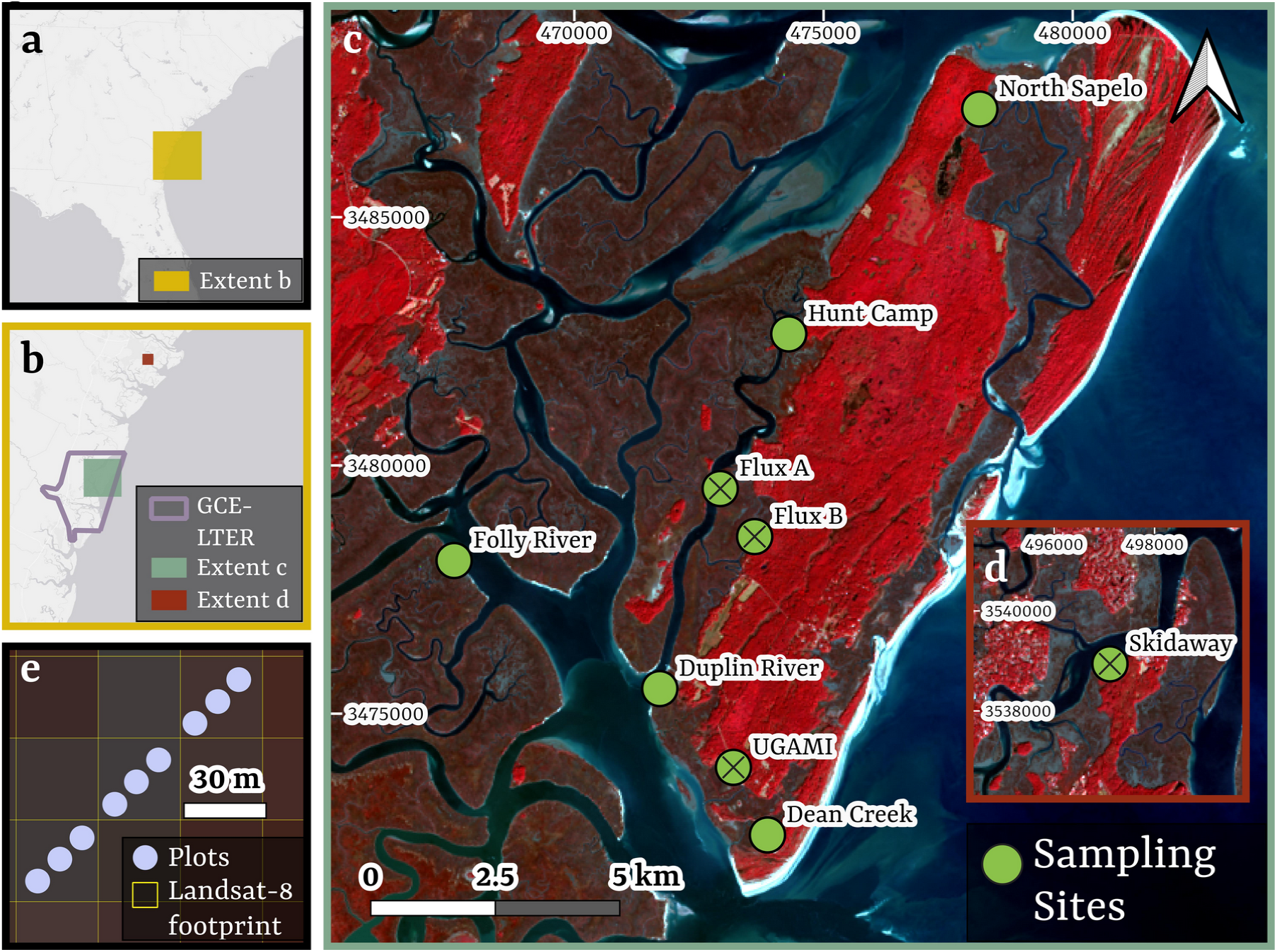
Field data were collected in *Spartina alterniflora* marshes in coastal Georgia, USA. a) Overview map showing the southeastern USA. b) Zoomed in overview map showing the boundary of the GCE-LTER. c) Site locations near Sapelo Island, GA, and d) Skidaway Island, GA. Text refers to site names. Circles with a cross depict sampling sites that were also used in O’Connell et al. 2021. e) Depiction of transect established at each site, with plots co-located with Landsat-8 and −9 pixel footprints. UTM coordinates reflect zone 17N. Basemap of panels a and b courtesy of ESRI. Basemap of panels c through e is false-color (R5, G4, B3) Landsat-9 imagery from 2024-01-20.

To characterize *in-situ S. alterniflora* BGB and later expand the BERM calibration dataset, we continued sampling at the four sites established by O’Connell et al. (2021) for initial BERM development, and added an additional five sites in diverse *S. alterniflora* marshes along the GA coast (Figure 1c, 1d; Table 1). Several sampling sites were located along the Duplin River or Doboy Sound, which are estuarine environments that receive variable freshwater input from the Altamaha River (Sheldon and Burd 2014). Newly established sites expanded the model parameter space by capturing variables across additional hydrogeographic settings, salinity, elevation gradients, and marsh channel morphologies. Mean channel salinity characterizations by Wang et al. (2017) suggest that the range of salinity was about 33% greater, and the range of elevation (as measured in this study) was nearly five times greater in the additional sites than in the previously established sites (Table 1).

**Table 1.** Newly established sites were selected to represent distinct hydrogeomorphic settings and expand model parameter space.

| Site<sup>1</sup> | Drainage Basin | Elevation Range<sup>2</sup> | *S. alterniflora* Height Forms Present<sup>3</sup> |
| --- | --- | --- | --- |
| Skidaway | Skidaway River | 0.79 to 0.87 | M, T |
| North Sapelo* | McCloy Creek | 0.77 to 0.89 | S, M |
| Hunt Camp* | Duplin River | 0.54 to 0.63 | M, T |
| Flux A | Duplin River | 0.65 to 0.82 | S, M, T |
| Flux B | Barn Creek | 0.69 to 0.75 | M, T |
| Folly River* | Folly River | 0.89 to 0.91 | S, M |
| Duplin River* | Doboy Sound | 0.91 to 1.07 | M |
| UGAMI | Lighthouse Creek | 0.75 to 0.85 | M, T |
| Dean Creek* | Dean Creek | 0.69 to 0.90 | M, T |

<sup>1</sup>Rows with an asterisk represent new data collection sites, while those without were established by O'Connell et al. 2021 and the GCE-LTER.; <sup>2</sup>m NAVD88; <sup>3</sup>S = Short, M = Medium, T = Tall, based on frequency of stem heights < 50 cm, between 50 and 80 cm, and > 80 cm;

### 2.2. Field Dataset

Our first objective was to characterize spatiotemporal variation in observed *S. alterniflora* BGB. The dataset included eight sites sampled quarterly from 2021 to 2023 (Feb, May-June, Aug, Nov) for a total of eight sampling events. Three of these sites had been previously sampled by O’Connell et al. (2021) over one growing season (five data collection dates in 2016), and these observations were included in the analysis. We also included data from the long-term sampling of the GCE-LTER site (monthly to quarterly from 2013 through 2023).

Data collection used the same methods as described in O’Connell et al. (2021) to ensure consistency. In brief, we established permanent 1 m^2^ field plots co-located within Landsat-8 and −9 pixel footprints at each of the eight sites, which supported model scaling through remote sensing methods. Field plots were placed along a transect that spanned a marsh edge to interior elevation gradient, allowing us to capture biophysical responses across the typical range of environmental characteristics at each site. Transects contained nine plots (plot-level), with three plots each per Landsat pixel (pixel-level; 30 m pixel resolution; Figure 1e). Multiple plots per pixel provided better pixel characterization and diminished the impacts of outliers. The GCE-LTER long-term site (‘Flux A’) consisted of targeted placement of between 2 to 12 plots within short, medium, and tall height forms of *S. alterniflora*, which are typically found along a marsh interior to edge elevation gradient. LTER staff did not deliberately co-locate these plots with Landsat pixels, and pixel footprints for this site had a variable number of plots.

Within our permanent vegetation plots, we used non-destructive methods to measure AGB ( g m^-2^;assessed allometrically from stem heights), leaf area index (LAI; unitless; assessed via AccuPAR sensor), and foliar chlorophyll (CHL; mg g^-1^; assessed via SPAD sensor calibrated with leaf chlorophyll extractions). See Appendix S1: Section S1.1., Appendix S1: Figure S1, and O’Connell et al. (2021) for a broad description of the methods. Aboveground leaf material from the root core (described below) was also processed (washed, dried, ground, and weighed) to measure foliar nitrogen with a CHN analyzer (N; %). Total foliar N (g m^-2^) was calculated by multiplying predicted foliar N (%) by predicted AGB (g m^-2^).

Belowground biomass was measured during each sampling event through the standing crop method (Graham and Mendelssohn 2016), where samples were collected via a standing root core adjacent to permanent plots. Root cores (7.62 x 30 cm) were centered on a random plant stem clump for collection (*n* = 1/plot/sampling event, totaling 749 novel cores for this study and 1099 total cores in combined calibration datasets with O’Connell et al. (2021); Appendix S1: Table S2). We also collected associated AGB present within the core footprint. Core above and belowground material was washed and sorted to retain live biomass (necromass or dead biomass was discarded), and then dried and weighed to develop a root:shoot ratio. This ratio was used along with allometrically assessed plot-level AGB to estimate plot-scale BGB.

Elevation at each plot was measured once in the NAVD88 vertical datum with a real-time kinematic global navigation satellite system RTK-GNSS (Trimble TSC7 handheld controller and R12 GNSS receiver; Trimble Inc., Westminster, Colorado, USA), and elevation measurements were related to the vertically benchmarked NOAA Fort Pulaski tide station (ID: 8670870).

### 2.3. In-Situ BGB Spatiotemporal Variation

We evaluated the annual cycle of BGB by grouping observed BGB by quarterly sampling event (i.e. Feb, May-Jun, Aug, and Nov), and visualizing through a loess regression. To test if BGB varied seasonally (i.e. differed among sampling events), we conducted Kruskal-Wallis tests by site. To evaluate the relative importance of spatial and temporal variation, we grouped and summarized field data categorically by site (site-level) and year (year-level) and used pairwise Wilcoxon rank sum tests (p < 0.05) to compare them.

### 2.4. Assembling a Candidate Predictor Dataset and Selecting Influential Predictors

We assembled a dataset of environmental variables as candidates to be applied as predictors of BGB in BERM. These candidate predictors included remotely sensed estimates of aboveground vegetation properties, elevation, tide, and climate data, which were grouped into categories of “biological,” “climatic,” “elevation”, and “hydrologic.” Highly correlated predictors were not excluded, as the decision tree-based algorithms used in our modeling framework (Appendix S1: Section S1.2.) were robust to multicollinearity (Mousa et al. 2019). Datasets of BGB and candidate predictors were spatially grouped and summarized by pixel footprint, and temporally matched to a monthly timestep through averaging and linear interpolation.

#### 2.4.1. Biological

We used the BERM workflow (Appendix S1: Section S1.2.) to predict field observed AGB, CHL, foliar N, and LAI from cloud- and tide-filtered Landsat-8 and −9 observations and associated indices (Appendix S1: Section S1.2.1., Appendix S1: Table S3). This involved building calibration datasets by linearly-interpolating the quarterly ground-truth dataset to a weekly timestep, where field observations within the same pixel were first averaged together.

This method took advantage of strategically timed collections of the ground-truth dataset, where we measured marsh attributes at key seasonal stages and assumed progressive growth between collection dates (O’Connell et al. 2021). The pixel-scale interpolated data was then temporally matched to the closest remote sensing observation. Prediction models were then built for each of the four aboveground plant biophysical variables measured (Appendix S1: Section S1.2.2.). Once aboveground models were created, predictions were made for each aboveground variable on all dates in the interpolated field dataset. NDVI, which responds to vegetation biomass and health, (Rouse et al. 1974, Glenn et al. 2008) was also included as a candidate predictor of BGB.

#### 2.4.2. Climatic

Local land surface temperature (LST), precipitation, solar radiation, and vapor pressure were obtained from Daymet, a set of daily 1 km gridded weather and climate estimates from ground-based observations and terrestrial modeling (Thornton et al. 2022). We also gathered 30 m LST from Landsat-8 and −9. We used the method of O’Connell et al. (2020) to estimate the day of *S. alterniflora* canopy green-up based on a growing degree day model of soil temperature. Soil temperature and growing degree day models were generalized to the site scale through Landsat LST. Annual day-of-year (DoY) of green-up, calculated through a growing degree days approach, can help anchor progressive seasonal productivity to the start of the growing season, which exhibits considerable spatial variation in this system (O’Connell et al. 2020).

#### 2.4.3. Elevation

Elevation, collected as a part of the ground-truth dataset (Section 2.2.), was the only candidate predictor considered in the self-titled category. Elevation was considered constant through time.

#### 2.4.4. Hydrologic

Hydrologic data were derived from remote sensing and tide data, and included three metrics for each pixel: flooding frequency (FF), inundation intensity (II), and dry intensity (DI). FF was the proportion of available remote sensing observations from 2016 through 2023 that were classified as flooded for that pixel, as estimated from a random forest remote sensing-based flooding model (described by O’Connell et al. 2021). Each pixel was associated with a single FF that did not change through time. II and DI were calculated monthly using FF, elevation, and tide data (monthly mean higher high and mean lower low water (MHHW, MLLW), respectively; Equations 1 and 2). Tide data were obtained from the closest NOAA CO-OPS benchmarked station, Fort Pulaski (ID: 8670870), located between 13 and 80 km northeast of our field sites. ElevationPixel was the pixel elevation.

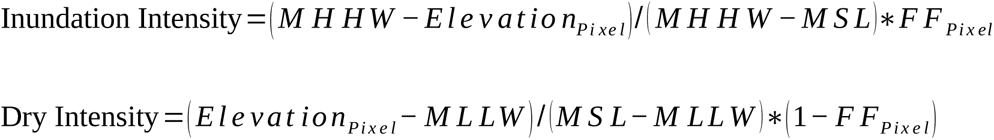

#### 2.4.5. Selecting Candidate Predictors for BERM

Candidate predictors of BGB for inclusion in BERM had to meet two criteria. First, we conducted an initial screening to determine if the variation in the candidate was related to the variation in observed BGB. This involved visualizing trends of candidate predictors against observed BGB through loess regression to explore the data and identify patterns. We retained only candidate predictors which co-varied with observed BGB. Pearson correlation coefficients between candidate predictors were assessed to broadly identify potential unique contributions to BGB predictions. Second, we used the BERM model training process (Section 2.5.) to identify and rank influential predictors of BGB. Here, we calculated time-series lags, rolling means, and differences of one through five months previous for each candidate predictor as described in O’Connell et al. (2021), because observed BGB may be a product of antecedent and longer-term conditions. Model feature importance scores were calculated based on the relative contribution to performance for each predictor variable, which indicates how valuable each predictor was for accurately predicting the response variable. Feature importance is a function of the contribution to model gain, coverage among observations, and frequency of feature use in trees and calculated in the ‘xgboost’ package in R (Chen et al. 2019). Candidate predictors with a model feature importance score below 0.005 during model tuning of the training set were discarded, and those remaining were retained and applied in BERM to estimate BGB (Appendix S1: Section S1.2.).

### 2.5. Advancing BERM

After identifying relevant candidate predictors, we advanced BERM from version 1.0 (BERMv1.0), which was the model described in O’Connell et al. (2021), to version 2.0 (BERMv2.0). Models were rebuilt using a framework similar to O’Connell et al. (2021) (Appendix S1: Figure S2, Appendix S1: Section S1.2.), using the identified predictors and the expanded ground-truth calibration dataset (Section 2.2). Critically, model training and testing were accomplished through spatiotemporal nested cross-validation to ensure that the model efficacy was evaluated against novel data not used to train the model. That is, data from each unique site and date group were blocked together while creating the training and testing splits as a measure to prevent overfitting, and all model tuning, feature selection, and fitting relied only on the training dataset, while final model performance was only evaluated against the testing set. Model performance was determined by mean bias error (MBE), root mean square error (RMSE), normalized RMSE (nRMSE), CoV RMSE (coefficient of variation of RMSE), and Pearson’s correlation of the field-observed vs. BERM-predicted testing data (for equations, see Appendix S1: Section S1.3.). Model predictions in testing splits among outer model folds (see Appendix S1: Section S1.2.) were averaged and plotted against observed BGB to assess model fit and bias. We identified thresholds for model under- and overpredictions by creating a linear model of the testing split residuals against the observations and identifying a BGB threshold where the residual as predicted by the linear model exceeded the BERM RMSE (Appendix S2: Section S2.3.).

We applied BERMv2.0 to demonstrate model utility and visualize marsh conditions at the GCE-LTER Flux Tower Marsh Sapelo Island, GA, in 2022 (Appendix S1: Section S1.4.). The Flux Tower Marsh consists of 1.3 km^2^ of *S. alterniflora* monoculture along the Duplin River and Barn Creek and has been extensively studied (Jung and Burd 2017, Tao et al. 2018, Alber and O’Connell 2019, O’Connell et al. 2021, Narron et al. 2022, Hawman et al. 2023, Hawman et al. 2024). We visualized BGB quarterly in 2022, and also produced marsh-wide estimates of AGB, LAI, CHL, foliar N, elevation, flooding frequency, II, and DI for May 15, 2022 as an example of BERM model output and data generation potential. A coast-wide application and quantitative assessment of BGB stock and trends is forthcoming in a future effort.

### 2.6. Spatial and Temporal Variation in BGB

We tested whether the BGB spatiotemporal dynamics observed in the field data were reflected in BERM output. To do this, first we compared variation in observed BGB among sites and years in field data by calculating the mean observed BGB by site and year, as well as the standard deviation of these means. A larger BGB range or standard deviation by site or year represented more dissimilarity. We also conducted a regression commonality analysis (CA) on pixel-level observed BGB to decompose explained variation of observations into unique and common effects among sites and years through a multiple regression analysis using the R package ‘yhat’ (Nimon et al. 2008, Nimon et al. 2021).

We applied similar methods to assess spatiotemporal dynamics as discerned through BERM, but used model error of the testing data (nRMSE) to compare means and standard deviation. We chose model error as our value of comparison instead of model predictions because error encompasses both observations and predictions, and helped to detect any biases that were present in the spatiotemporal structure between the two. Here, the minimum number of testing data observations was set to 10. Each of the nine sites and the eight years present in the calibration data then were associated with a set of error metrics. Group means were compared using Wilcoxon rank sum tests. A larger mean or standard deviation nRMSE represented poorer model performance, and can be associated with a larger degree of observed variation in that group given the assumption of a generalized model. We conducted a CA on model predictions with the same effects as done with observed BGB.

## 3. Results

### 3.1. Observed Biomass Dynamics

Observed BGB varied across both space and time (Appendix S1: Table S4). The overall mean observed BGB was 989 g m^-2^ with a high standard deviation (912 g m^-2^; Appendix S1: Figure S3a) and variation among sites (Appendix S1: Figure S3b). We observed a weak seasonal trend in BGB (Figure 2). BGB was highest in February, then fell in May-June and August, and slightly rose in November. Only at two sites, Folly River and UGAMI, was BGB different among sampling events, indicating a significant trend. Overall, differences among sites were greater than among years (Figure 3). In 69% of pairwise comparisons, distributions of observed BGB were significantly different by site, and no sites were universally similar. By year, observed BGB distribution significantly differed in only 21% of cases, and these significant comparisons all involved the years 2017 and 2018. Observed BGB distributions in these years were more bimodal and skewed-right; that is, they had more high BGB observations.

**Figure 2.**
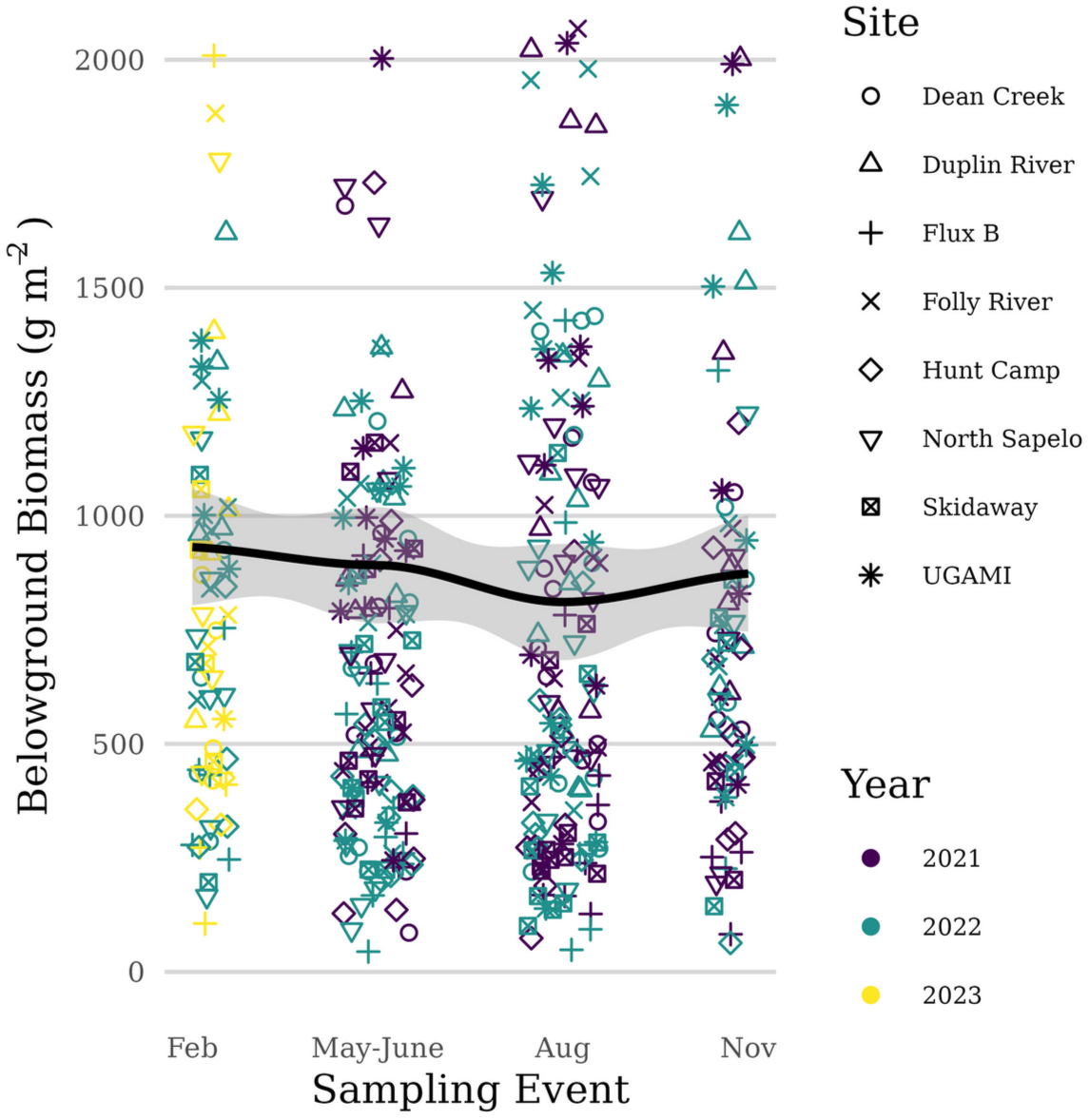
Plot-level seasonal belowground biomass patterns. The black line depicts a loess regression. Outliers above 2000 g m^-2^ were removed for visualization (*n* = 36). Data presented consists only of new data collection for this study for consistent seasonal sampling.

**Figure 3.**
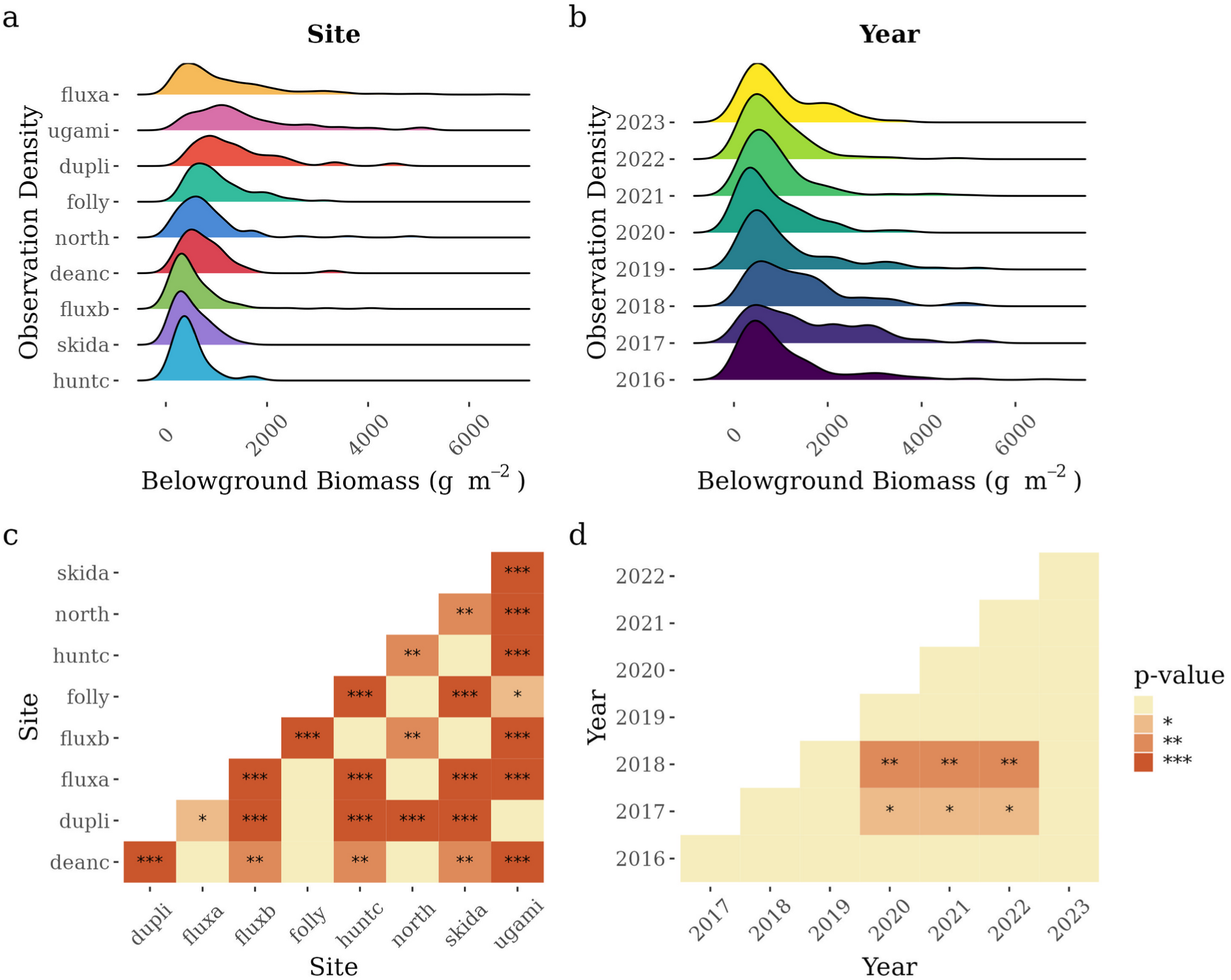
Comparisons of site and year level observed belowground biomass (BGB). Density plot of observed BGB by site (a) and year (b). Pairwise wilcox-test significance matrix of levels within site (c) and year (d) groupings. Blank: not significant; *: p < 0.05; **: p < 0.01; ***: p < 0.001.

### 3.2. Candidate Predictors

Environmental variables selected as candidate predictors exhibited varying trends against observed BGB (Figure 4). Trends of biological candidate predictors against observed BGB typically displayed roughly unimodal response curves. In other words, maximum observed BGB was found at intermediate values of each biological candidate predictor. The root:shoot ratio (AGB:BGB) was highly variable (mean: 4.3, standard deviation: 3.5). Candidate predictors in the climatic category did not appear to be strongly related to observed BGB, as indicated by mostly flat relationships, with the exception of those related to temperature. The highest values of LST were associated with higher BGB. Green-up DoY also had a potential positive trend with BGB at later dates. In terms of hydrologic variables, BGB peaked at low FF and II and then decreased, whereas DI showed the opposite pattern, again peaking at a local maximum. Elevation displayed a distinct trend against observed BGB. An optimal elevation for BGB existed at about 0.8 m NAVD88 (equivalent to 0.9 m at the MSL datum, −0.1 m at MHW, and −0.2 m at MHHW, relative tidal elevation, Z*, of 0.9, as defined in Morris et al. (2002) and Swanson et al. (2014)). Most sites (five of nine) spanned this optimal elevation, while two were entirely below this elevation, and two entirely above (Appendix S1: Figure S4). While BGB was maximum at this elevation, the relationship otherwise appeared similar to a step function, where low BGB was found at elevations below 0.7 m NAVD88 and high BGB at elevations above 0.9 m NAVD88.

**Figure 4.**
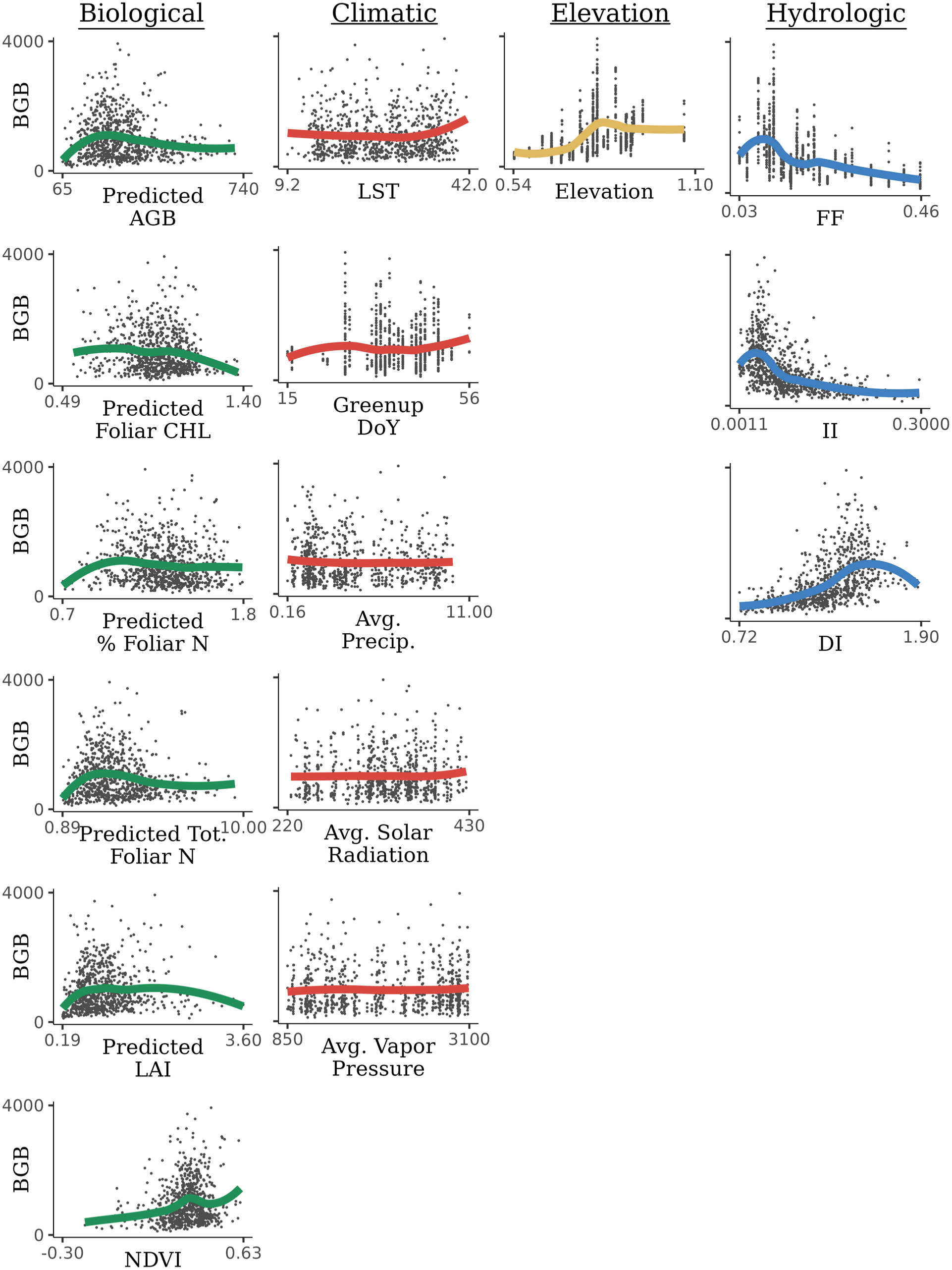
Candidate predictor trends with pixel-level observed belowground biomass (BGB). In each plot, the candidate predictor is the independent (x) variable, and observed BGB is the dependent (y) variable. Units for candidate predictors are as follows: Predicted aboveground biomass (AGB; g m^-2^), Predicted % Foliar Chlorophyll (CHL; mg g^-1^), Predicted % Foliar Nitrogen (N; %), Predicted Tot. Foliar N (g m^-2^), Predicted Leaf Area Index (LAI; unitless), NDVI (unitless), Land Surface Temperature (LST; degrees C), Greenup Day of Year (DoY), Avg. Precip. (mm day^-1^), Avg. Solar Radiation (W m^-2^), Avg. Vapor Pressure (Pascal), Elevation (m NAVD88), Flooding Frequency (FI; proportion), Inundation Intensity (II; proportion), and Dry Intensity (DI; proportion). The scatterplot represents observed data. The curves, color coded by category, represent loess regressions. Candidate predictors that show distinct trends with BGB may be influential in BGB prediction.

Influential predictors of BGB were identified through ranked feature importance scores from the BGB models (Figure 5). We removed three potential predictors from the modeling framework following importance and co-variance screening. These three (precipitation, solar radiation, and vapor pressure) each belonged to the climate category. All other candidate predictors considered covaried with BGB (Figure 4) and were found to have a feature importance above the 0.005 cut-off (either coincident or in a time-integrated form; Figure 5). Most variables retained were a rolling mean, lag, or time-series difference of on-the-ground characteristics, rather than current values for the date of prediction.

**Figure 5.**
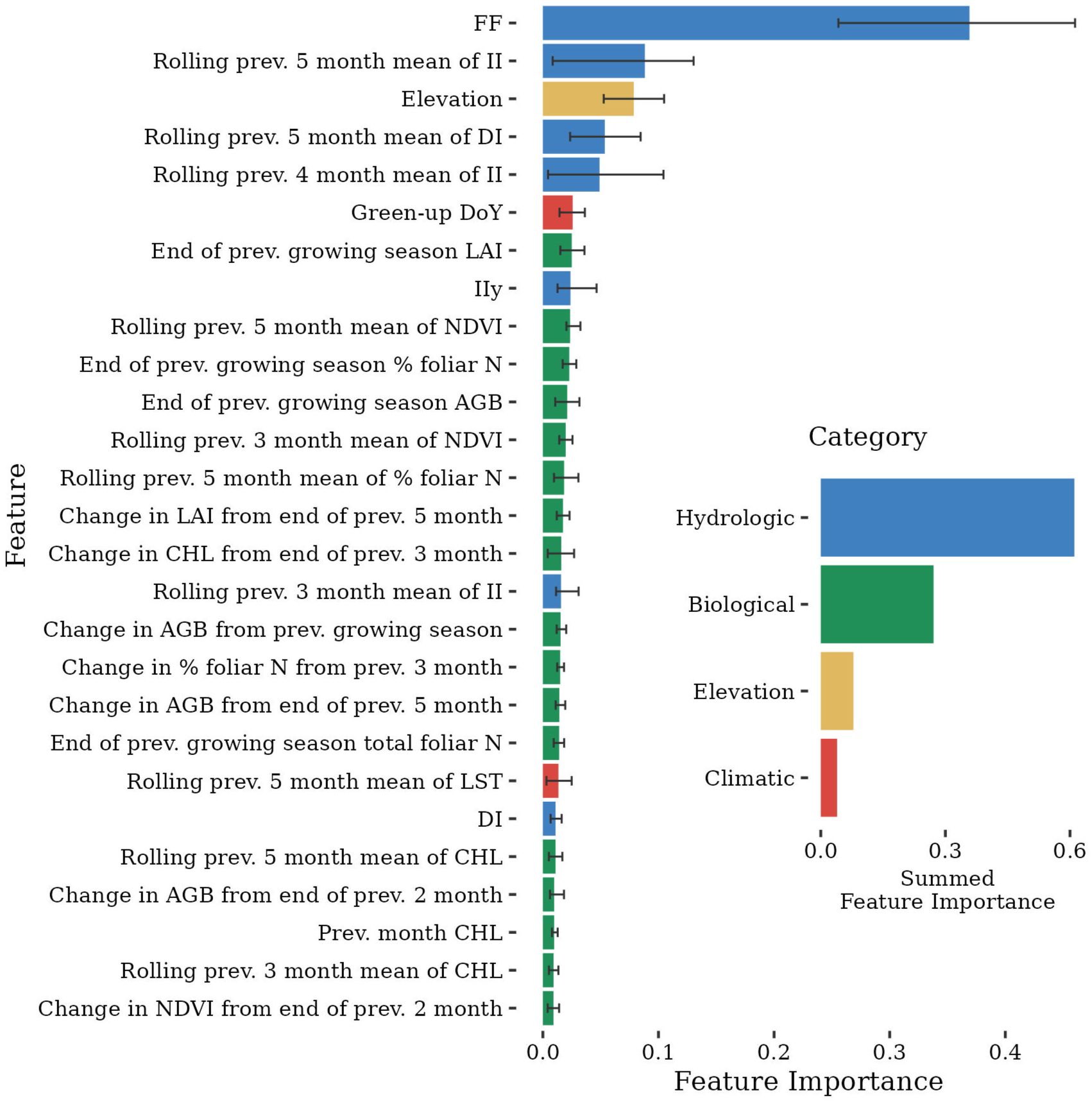
Main plot: Average feature importance of the 27 features selected for inclusion in the final belowground biomass model. Error bars depict the range of importance among the five model outer folds. Features are color-coded by categories Biological, Climatic, Elevation, and Hydrologic. Abbreviations: FF: Flooding Frequency; II: Inundation Intensity; DI: Dry Intensity; DoY: Day of Year; LAI: Leaf Area Index; NDVI: Normalized Difference Vegetation Index; AGB: Aboveground Biomass; CHL: Chlorophyll; LST: Land Surface Temperature; N: Foliar Nitrogen. Inset plot: Sum of the feature importance by category.

Flooding frequency was by far the most important predictor variable, with a feature importance of 37%. Flooding, combined with other hydrologic variables had the largest sum of feature importance (61%), followed by biological (27%), elevation (8%), and finally climatic (4%) related variables (Figure 4). Most of the summed feature importance of all predictors (64%) was accounted for by just five variables.

Few candidate predictors were strongly correlated with each other. Overall, only 11% of relationships between candidate predictors resulted in a correlation coefficient above 0.5 or below −0.5, and most of these involved related variables (i.e., a variable used to calculate another). This typically occurred within categories (biological: 25% of relationships, climatic: 22%, elevation: NA, hydrologic: 100%). FF and elevation were moderately correlated (r = - 0.32).

### 3.3. Advancement to BERMv2.0

We advanced BERM to v2.0 with the expanded calibration dataset and updated candidate predictors. The expanded calibration dataset broadened the range and altered the shape of observations for all vegetation metrics (above- and belowground; Appendix S1: Figure S5). Performance metrics of the updated model indicate good fit and low model error (Figure 6, Table 2). Compared to the v1.0 model, the RMSE and nRMSE decreased by 15% and 28%, respectively (Appendix S1: Section S2.2.), reflecting an improvement in performance.

**Figure 6.**
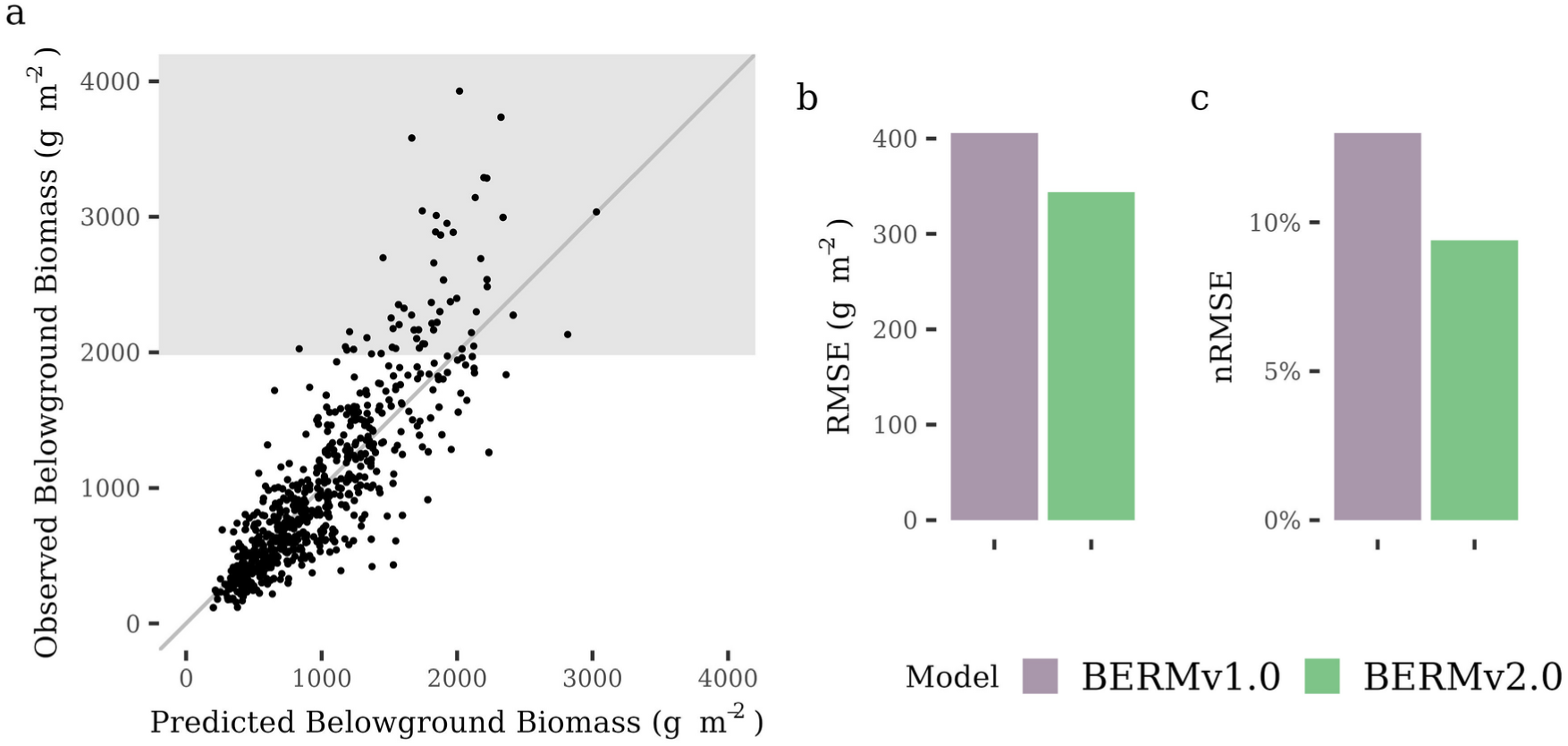
a) Observed vs. predicted BGB in BERMv2.0. Predicted BGB represents an average of the five model outer folds for the testing splits only (Methods S2). BERMv1.0 versus BERMv2.0. The grey shaded area represents observed BGB greater than 1981 g m^-2^, the threshold which we identified above which BERM underpredicted BGB. b) Root Mean Square Error (RMSE) and c) Normalized Root Mean Square Error (nRMSE).

**Table 2.** Testing data mean goodness of fit metrics for the *Spartina alterniflora* belowground vegetation Extreme Gradient Boosting models.

| Model | MBE<sup>1a</sup> | RMSE<sup>1b</sup> | nRMSE<sup>c</sup> | COV<br>RMSE<sup>d</sup> | Correlation | *n*-Test | *n*-Train |
| --- | --- | --- | --- | --- | --- | --- | --- |
| BERMv1.0 | -25.01 | 405.80 | 13.0% | 31.9% | 0.86 | 53 | 118 |
| BERMv2.0 | -5.03 | 343.76 | 9.4% | 35.6% | 0.85 | 225 | 531 |

<sup>1</sup>g m<sup>-2</sup>; <sup>a</sup>Mean Bias Error; <sup>b</sup>Root Mean Square Error; <sup>c</sup>Normalized Root Mean Square Error;
<sup>d</sup>Coefficient of Variation of Root Mean Square Error;

We evaluated the tendency of BERM to under or overpredict the observed data. A threshold for underprediction (residuals exceeded RMSE via a linear model with observations) was found above 1981 g m^-2^, and no threshold for overprediction was found (Figure 6; Appendix S1: Figure S6). This was mostly driven at one ‘Flux A’ pixel in short-form *S. alterniflora*, where the pixel MBE was −118.4 g m^-2^. BERM produced underpredictions only slightly more often than overpredictions at this pixel (32% and 23% of predictions, respectively). Further, large underpredictions (e.g. those where the individual prediction was less than the observation minus three times the RMSE) were rare (9% of predictions), while no large overpredictions were made. In these rare large underpredictions, the ratio of the stem densities within the root core footprint and the nearby vegetation plot drastically differed from the overall average. Our process to scale-up core-level BGB to plot-level BGB relied on root:shoot ratio and stem counts within both the root core and vegetation plot. Thus, it is possible that our upscaling process misrepresented observed BGB in the uncommon instances where core- and plot-level stem densities were very different.

To demonstrate BERM, we provide example output for the GCE-LTER flux tower site in Figure 7. The patterns of BGB over the course of a year show evidence for both intra-annual variation and spatial variability. In addition to predictions of BGB, BERM also produces spatially-explicit information on additional metrics which may be useful in marsh assessments. In this example one can see that the higher BGB values observed in May 2022 are often in well-drained areas of the marsh interior (e.g., areas with higher DI). Some areas show persistent low BGB, especially in concert with high FF. Notably, AGB patterns do not mirror BGB, suggesting the need for separate models of each parameter.

**Figure 7.**
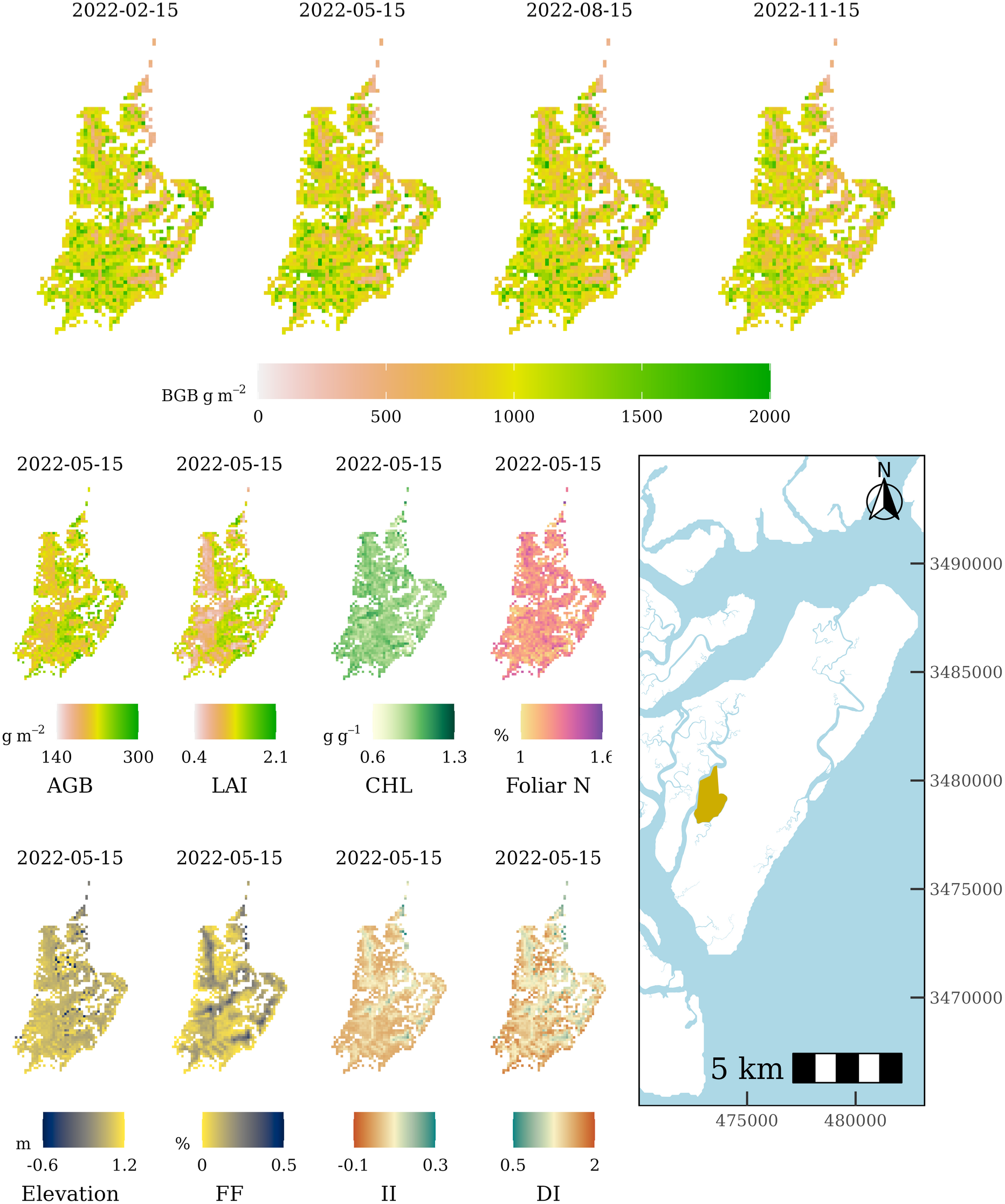
BERM application demonstration showing belowground biomass (BGB) estimated quarterly during 2022. Aboveground Biomass (AGB), Leaf Area Index (LAI), Foliar Chlorophyll (CHL), Foliar Nitrogen (N), and slower changing variables, such as elevation (datum: NAVD88), flooding frequency, inundation intensity, and dry intensity were estimated or calculated in May 2022. The Flux Tower Marsh on Sapelo Island, GA, USA is shown in gold in the lower right; coordinates refer to UTM zone 17N.

### 3.4. Observed and Model-Predicted Spatiotemporal Dynamics of BGB

BERM predictions reflected observed BGB spatiotemporal dynamics. The range of mean observed BGB was 26% greater by site than by year, and the mean model error was 51% greater (Figure 8a, 8c). Standard deviation of these means were also greater among spatial than among temporal distributions (observations: 38%; model error: 55%). In both observed and model predicted BGB, a low total amount of variation in BGB was explained through variance partitioning (observations: 33%; model predicted: 34%), but majorities of explained variance could be attributed to the spatial component (observations: 84%; model predictions: 85%; Figure 8b, 8d; Appendix S1: Table S6). Each of these comparisons show that the spatial component dominated the signal of observed BGB, which was also reflected in model outcomes.

**Figure 8.**
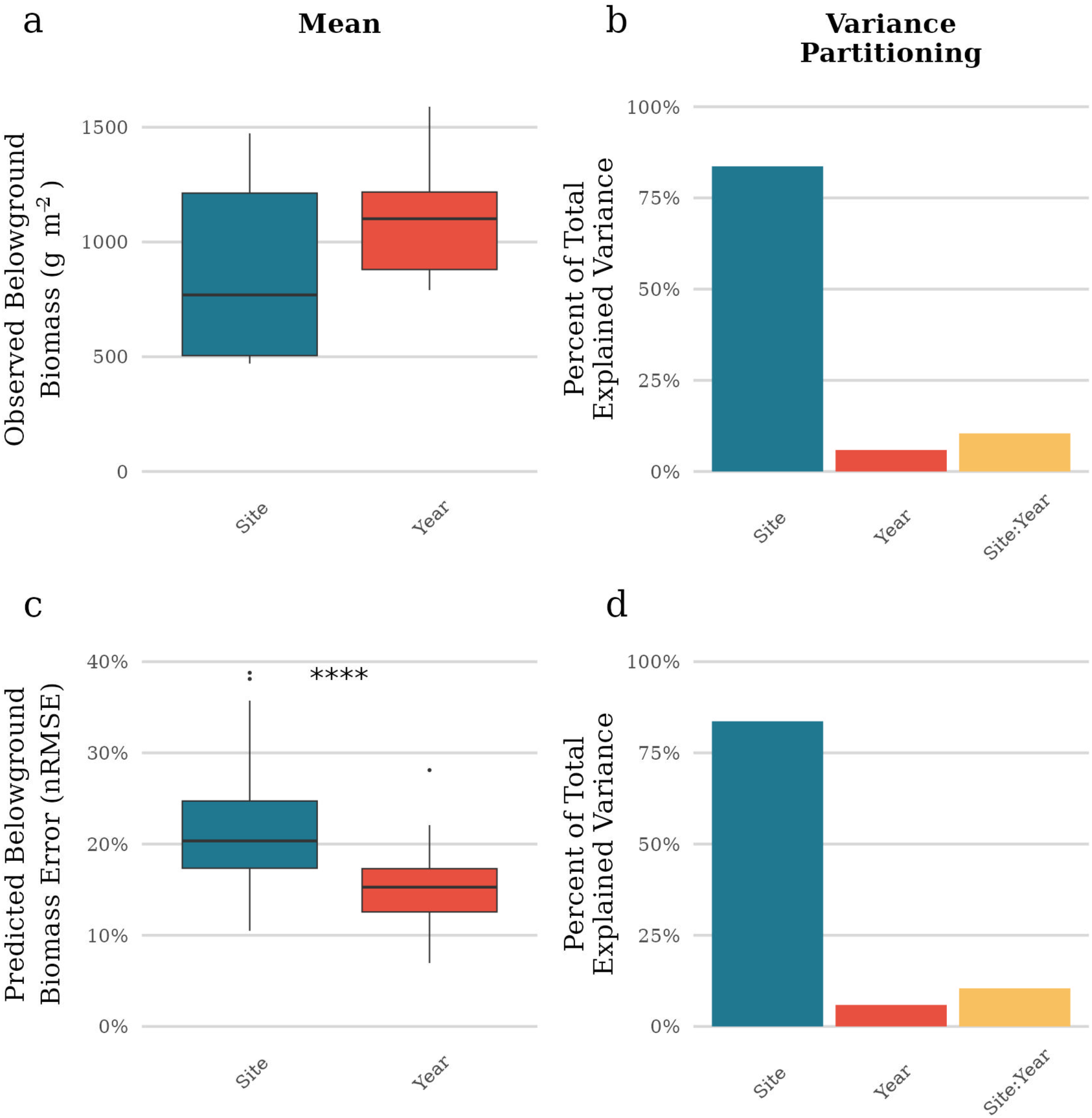
a) Mean observed BGB, grouped by site and year. (b) Variance partitioning through regression commonality analysis of observed BGB with components of site, year, and a site:year interaction. (c) Mean error (nRMSE) of predicted BGB, grouped by site and year, ***: p < 0.0001. (d) Variance partitioning through regression commonality analysis of predicted BGB with components of site, year, and a site:year interaction.

## 4. Discussion

Our analysis and update of BERM to v2.0 took advantage of one of the longest continuous BGB data collections through the GCE-LTER, and complemented it with a comprehensive set of additional *S. alterniflora* sites that represent conditions within GA marshes. A dataset of this size for *in-situ* marsh BGB is difficult to find outside the scope of this work. This allowed us to extend and improve BERM, a geospatial tool that can be used to provide new insight into marsh BGB dynamics and its potential as a key indicator of marsh resilience.

### 4.1. Field Patterns of BGB

The dataset we have compiled here is one of the most comprehensive of its kind to quantify salt marsh BGB across a region over multiple years, and has shown the relative contributions of broad components of variation. The high spatiotemporal variability of *S. alterniflora* AGB has been well-documented (Gross et al. 1990, O’Donnell and Schalles 2016, Eon et al. 2019, Zhu et al. 2019), but that of BGB has limited examples. Descriptions of *in-situ* spatiotemporal variation of *S. alterniflora* BGB stock in the literature rarely have focused on annual variation; most accounts captured seasonal changes, or intra- and inter-site variation (Ellison et al. 1986, Gross et al. 1991, Connor and Chmura 2000). Strong seasonal patterns have been theorized in BGB (Gallagher 1983), which we similarly observed, albeit weakly (Figure 2). Long-term trends are hard to capture with field data alone, given the difficulty of measuring BGB and the high variability it exhibits.

In this study we found that BGB was more spatially than temporally variable. In other words, BGB varied more across different locations of the landscape than it did between years. This is an important consideration for sampling design when researchers want to understand salt marsh BGB stock. Our finding that spatial gradients dominate the variation signal demonstrates that, given limited resources, ground-truth data collection campaigns may be more effective by prioritizing sampling at diverse sites at the expense of multi-year sampling. Yet, interannual variation can be substantial: pairwise comparisons did show that two years significantly differed from others, as 2017 and 2018 had more high BGB observations (Figure 3). A number of broad-scale phenomena could drive interannual variation, such as episodic storm events, variable freshwater inflow, and climate (Alber et al. 2008, Walters and Kirwan 2016, Alldred et al. 2017). For instance, hurricane Matthew caused extensive dune overwash along the GA coast in October 2016 (Birchler et al. 2019), potentially nourishing the salt marshes with sediment and stimulating belowground growth in the following years (Elsey-Quirk and Unger 2018), which may have explained our observed high BGB during 2017 and 2018.

### 4.2. Co-Varying Environmental Variables are Influential Predictors of BGB

We found that candidate predictors related to biological, climatic, hydrologic, and physical conditions varied with BGB and were influential predictors in our modeling framework. Most candidate predictors exhibited a non-linear relationship with observed BGB (Figure 4). This highlights the need for a wide range of calibration data in developing accurate prediction models, as non-linear relationships can appear linear over short ranges. Few candidate predictors were strongly correlated with one another, particularly between categories, implying broad suitability in providing unique information to estimate BGB.

Hydrologic metrics were the most influential predictors of BGB, and flooding frequency was by far the single most important metric (Figure 5). Studies have focused on this relationship with marsh organ mesocosm experiments (as introduced by Morris (2007)); for example, Snedden et al. (2015) and Hanson et al. (2016), in microtidal conditions, found that *S. alterniflora* BGB decreased with increased flooding duration. These studies assessed this relationship through six and three flooding frequencies (defined by target elevations), respectively. These frequencies covered a large extent (3.1% to 87.8% time inundated), but then also had wide ranges between treatments, and thus were unable to resolve subtle responses. On the contrary, we found a peak BGB at low to intermediate levels of flooding (∼10% flooding frequency). Our *in-situ* study, with BGB observations across 32 different flooding frequencies, provided higher resolution insight into this relationship and suggests that fine-scale variation results in roughly unimodal response curves rather than a linear or exponential fit.

Elevation may often be considered as a proxy of flooding, but we found it important to uncouple elevation and flooding frequency. Marsh flooding is a function of complex geomorphology, where not only salt marsh elevation, but also differences in tidal creek branching patterns, can admit or restrain tidal water delivery and draining (Friedrichs and Perry 2001). This has led to a recognition that general bathtub models, which use tidal elevation to predict flooding frequency, often cannot resolve marsh flooding patterns (Fagherazzi et al. 2012, O’Connell et al. 2017). Our finding that elevation and flooding frequency were only moderately correlated supports this point, and that flooding frequency was a stronger predictor of BGB highlights how important this distinction is. We estimated flooding through a remote sensing-derived pixel classification, which is unique from DEM-derived estimates of elevation. This classification likely better depicted hydrodynamics across the heterogeneous salt marsh landscape which are collectively influenced not only by creek branching patterns, but also complex microtopography, migrating tidal creeks, wrack deposition, wind dynamics, and accretion/subsidence processes. These diverse landscape features act together to alter marsh geomorphology and drainage on short spatial and temporal scales (Vu et al. 2017, Craft 2023, Lynn et al. 2023).

Our aboveground biological metrics, which mostly consisted of AGB and leaf biomass composition, also varied with BGB; typically, intermediate values of AGB were associated with the greatest BGB. But, these relationships were noisy, and did not rank high in feature importance in our modeling framework. Further, we found high variation in root:shoot ratio, indicating that AGB stock does not reliably indicate BGB stock. While the amount and composition of AGB have been designated as signals of marsh health (Marsh et al. 2016), assessments based solely on aboveground metrics may not provide a complete picture as biomass allocation (root:shoot ratio) varies over environmental gradients (Darby and Turner 2008a, 2008b, O’Connell et al. 2015, Li et al. 2021). Other factors likely affected aboveground plant growth that we did not directly measure, such as freshwater delivery and herbivory (O’Donnell and Schalles 2016, Wittyngham 2021, Niu et al. 2023). However, responses to these factors were captured by the aboveground biophysical variables that we did measure, allowing BERM to estimate complex interactions between BGB and the environment.

The relationship between BGB and temperature can be difficult to disentangle from seasonal growth patterns and phenological adaptation, as few studies have examined BGB across multiple years. Temperature may impact BGB growth and decomposition, as Crosby et al. (2017) showed that higher latitude, lower temperature marshes had greater BGB stock than those at lower latitudes and higher temperatures. But, this could be a phenological adaptation, and long-term warming may be stressful to plants (Crosby et al. 2017). The timing of changes in temperature may be impactful, too, as O’Connell et al. (2020) theorized that warm winter temperature could lead to high respiration and overwintering BGB depletion, perhaps representing an under-recognized climate vulnerability. In support of this theory, we observed low BGB in years of very early green-up (∼20% below average), though beyond that, the relationship did not appear to be strong. However, the set of environmental conditions BERM provides invites investigation of complex undiscovered relationships and multiple ecological stressors in future studies.

Contemporaneous and antecedent environmental conditions were influential in estimating *S. alterniflora* BGB stock. Metrics up to five months previous, as well as those from the end of the previous growing season, were selected in our modeling framework. Such preceding conditions may be important because of seasonal trends, observed directly in climatic predictors, and indirectly in others. For example, the rolling mean of the previous 5 months of LST was identified as an important predictor of BGB. Broadly speaking, in summer or winter, the preceding 5 month average temperature would be an intermediate value, whereas in spring or fall, it would be low or high, respectively. The trend of LST showed highest BGB at low or high temperatures, so the modeling framework may have related an intermediate preceding average LST with high winter BGB. The prevalence of these lagged and time-integrated features may also indicate that BGB is slow to change, aside from state transitions, which have been observed at our study location (Alber et al. 2008, Elmer et al. 2013). Roots exhibit longevity and turnover time in *S. alterniflora* and related species have been found to be slow, on the scale of months to years (Smith et al. 1979, Blum 1993, Bouma et al. 2003). These findings and characteristics of *S. alterniflora* root biomass suggest some resilience to short-term stressors.

By observing a wider range of phenomena, and subsequently updating our ecosystem prediction tools, we have refined how we understand ecological interactions between our predictor and response variables. Features retained in BERMv2.0 varied slightly from those used in BERMv1.0, as reported by O’Connell et al. (2021). Both versions of the model included variables within all categories we identified as broadly important groups of conditions. But, certain variables deemed important by O’Connell et al. (2021) in BERMv1.0 were discarded in BERMv2.0, including the second and fifth most important features in BERMv1.0, vapor pressure and precipitation, respectively. The BERMv1.0 calibration dataset was limited to collections from May through October in 2016 at three of four sites. These months generally represent the highest vapor pressure and precipitation throughout the year in coastal GA. Thus, site-level differences in BGB may have been associated with high summer variation in vapor pressure and precipitation in BERMv1.0. Our updated BERMv2.0 calibration dataset had more thorough temporal coverage, where all sites were associated with at least two years of data collection and was less vulnerable to this potential proxy effect.

### 4.3. BERMv2.0 as an Enhanced Ecosystem Prediction Tool

The performance of BERMv2.0 was strong, as assessed by rigorous evaluation of the error through five metrics on novel testing data. We found that BGB varied more over spatial gradients than temporal (Section 3.1). Thus, by focusing on broadening the spatial calibration in building the v2.0 calibration dataset, we likely represented variation across the landscape more realistically. This helped reduce the BERMv2.0 error when it was applied across a wider parameter space. Products from BERMv2.0 include estimates of aboveground parameters and BGB, and can be used to describe resilience or support other marsh investigations.

BERM is the first of its kind BGB model that estimates spatiotemporal variation in near real-time, as informed by satellite and ground-based observations to estimate realistic BGB trends with spatial context. BERMv2.0 is archived on GitHub and available for use by the scientific community. The model was created in reproducible modeling workflows that rely on openly available and free data, and thus BERM application was designed to be easily deployable to generate detailed landscape assessments. Products such as maps of BGB change can guide land management such as conservation and restoration, and provide insight into drivers of BGB. The model workflow is also transferable to new regions and species, given sufficient calibration data and use of the methods described here. Built with Landsat-8 and −9, BERM is compatible with nationwide coastal wetland data products, but also could be calibrated in the future to utilize Sentinel-2, which provides finer spatial, spectral, and temporal resolution. Sentinel-2 particularly has red-edge bands which can be informative in estimating vegetation stress, chlorophyll, and foliar nitrogen.

### 4.4. Model-Building to Capture Key Components of Variation

Ecosystem models such as BERM aim to describe trends across a landscape, and trend detection capability depends on components of variation (Larsen et al. 2001). In our observed BGB, we found the spatial component to dominate the variation signal. Thus, in building a BGB prediction tool, we should strive to reflect this systematic variability. Model performance metrics such as MBE, RMSE, and nRMSE can indicate average prediction accuracy, but alone they do little to indicate spatial or temporal variation in model performance and predicted data. By showing that the spatiotemporal structure of observed BGB was matched by model performance, along with similar variance partitioning, we confirmed that BERM reflects the systematic variability of our target ecosystem.

## 5. Conclusion

Quantitative estimates of salt marsh BGB are critically important for understanding marsh resilience to sea level rise, but traditional field surveys are unable to appropriately capture BGB variation over space and time. Ecosystem models such as BERM can capture this variation, but calibration requires attention to characteristics of variation across the landscape to represent trends in productivity. We measured BGB in GA *S. alterniflora* marshes across an extensive range of environmental conditions. We found that spatial variation dominated the signal of BGB, as it varied more across short spatial scales than it did among consecutive years. Coincident environmental conditions, particularly those related to hydrology, emerged as strongly related candidate predictors of BGB. With this, we advanced BERM to version 2.0 as a geoinformatics tool to evaluate landscape scale change in U.S. GA *S. alterniflora* salt marshes.

The ability to estimate BGB across time and space will fill a critical gap in better understanding marsh plant productivity, wetland vertical accretion, and carbon dynamics (Osland et al. 2022, LaFond-Hudson and Sulman 2023) and can serve as a primary metric in prioritizing areas for coastal restoration. BERM estimates, based on empirical conditions, can not only be used in management decisions themselves, but can be coupled with process-based models such as the Cohort Marsh Equilibrium Model (Vahsen et al. 2024) to incorporate realistic BGB estimates across time and space to improve marsh accretion estimates. Additionally, BERM predictions can refine blue carbon assessments, as BGB estimates, paired alongside those of AGB, can directly be applied to quantify net primary productivity and carbon accounting (Woltz et al. 2023). Blue carbon dynamics can provide further insight into microbial ecology as leaky root exudates fuel microbial and higher trophic food webs (Turner 1993). The BERM framework also generates a suite of predictor metrics, which themselves represent useful information in salt marsh investigations. For example, estimates of foliar chlorophyll and nitrogen may illustrate coastal nutrient flow, while flooding and temperature could parameterize decomposition models. And ultimately, changing phenology of each of these metrics with global climate change can be captured through the near real-time estimates.

## Data availability

The complete ground-truth dataset is archived at the GCE-LTER online data portal: https://doi.org/10.6073/pasta/4a0b715104849d98320fcc34e7cd63a4

BERMv2.0 code is available at: https://github.com/kylerunion/berm_v2

## Author contributions

KDR, DMR, MA, and JLO planned and designed the research. KDR collected the field data, conducted the analysis, and wrote the manuscript. DMR, MA, MAL, and JLO contributed to writing the manuscript and interpreting the results.

## Acknowledgments

This work was supported by the National Oceanic and Atmospheric Administration Margaret A. Davidson Graduate Fellowship [grant number NA22NOS4200060], and the National Aeronautics and Space Administration Future Investigators in Earth and Space Science [award number 80NSSC23K0300]. The authors also received support from the GCE-LTER, which is supported by the National Science Foundation (OCE-1832178). This is UGAMI contribution number 1121. We thank Wade Sheldon, Adam Sapp, Dontrece Smith, John Williams, Jonah Rigdon, Karolena Popyk, Jackson Akin, Ryan Hladyniuk, and Patty Garlough for aiding in collection, processing, and archiving of marsh productivity data, and Kristin Byrd, for contributions to BERMv1.0 that were influential to this work.

