## Supplemental File for "Capturing spatiotemporal variation in salt marsh belowground biomass, a key resilience metric, through geoinformatics"

### Supporting Information

### Section S1: Methods

#### *S1.1. Chlorophyll extraction for SPAD meter calibration*

The Konica Minolta Chlorophyll Meter SPAD-502Plus (SPAD) estimates leaf chlorophyll by calculating the optical density difference of two wavelengths. LED light sources in the red and near-infrared regions of the electromagnetic spectrum shine light on the leaf to be measured, and absorbance is measured. We used two different SPADs for field chlorophyll measurements. Measurements may not be consistent among machines, so each machine should be calibrated against laboratory-derived chlorophyll measurements (Markwell et al. 1995). The calibration of the first SPAD meter, which measured chlorophyll for all samples before 2021, was described in O'Connell et al. (2021). This section describes the calibration process for the second SPAD meter, which was used for all chlorophyll measurements taken after May 2021.

To calibrate our SPAD, we followed the protocol established by Ni et al. (2009), modified to use ethanol as a solvent as described by Ritchie (2006). SPAD measurements of 44 leaves were taken in the field (3 replicates per leaf, each wiped clean with a wet rag to remove dirt particles and salt films) with a range of yellow and green leaves targeted. Each leaf was clipped and individually wrapped in aluminum foil to prevent light exposure, then stored on ice to minimize chlorophyll degradation before being transported to the laboratory. Chlorophyll extractions with 100% ethanol were completed within 12 hours of sample collection. Absorbance of the ethanol and chlorophyll solution was measured after an overnight extraction at 649 nm and 665 nm with a spectrophotometer. Leaf chlorophyll concentration (CHL) was calculated as:

$$CHL_a(\mu g/g)=[-5.2007*A_{649}+13.5275*A_{665})*V/W$$

$$CHL_b(\mu g/g)=[22.4327*A_{649}-7.0741*A_{665})*V/W$$

$$CHL=CHL_a+CHL_b$$

where A = absorbance at wavelength of subscript, V = volume of extraction (mL), W = weight of fresh leaves (g)

Once we had SPAD measurements and CHL for each sample, we compared linear and exponential regression to estimate leaf chlorophyll from SPAD measurements. The linear model explained 63% of the variation between SPAD measurements and leaf chlorophyll, and the exponential model explained 73% (Figure S1). Though the exponential model explained more variance, BERMv2.0 model performance did not differ between calibration data calculated with the linear and exponential model. For consistency with O'Connell et al. (2021), we calculated leaf chlorophyll concentration with the linear model.

The linear model equation was:

$$CHL(mg/g)=-0.32+0.038*SPAD$$

The exponential model equation was:

$$CHL(mg/g)=e^{0.06*SPAD+-2.454}$$

*S1.2. Updating BERM*

BERM consists of a suite of models to predict four aboveground metrics and belowground biomass (BGB). BERM models utilized extreme gradient boosting (XBG), a decision tree ensemble machine-learning algorithm, used here for a regression-based prediction model. We used the 'XGBoost' package in R (Chen et al. 2023) to build prediction models. To create and calibrate models, first, we divided calibration data into training and testing splits through nested spatiotemporal cross-validation. This was an important step to prevent biases and overconfidence in model results (Roberts et al. 2017, Schratz et al. 2019). Inner cross-validation was used for hyperparameter tuning, whereas outer cross-validation splits were reserved for final model validation. That is, testing data were not accessible to the model while developing relationships between predictors and the variable of interest during model tuning. As another check, we kept data from the same date and site together in either the training or testing sets. This avoids splitting highly spatiotemporally correlated data amongst the training and testing data, as that would result in over-optimistic model evaluation because the training and testing data would be more closely related than novel landscape data. Models of above- and belowground vegetation characteristics followed different model training frameworks to incorporate an appropriate amount of resources. Aboveground models were built and assessed with cross-validation of one outer fold and three inner folds, while the belowground model, which is more complex, applied five outer and five inner folds. Cross-validation with inner folds are important for assessing model performance on the training data to prevent overfitting, and with the outer folds for assessing performance on 'unseen' testing data. Overfitting is a phenomena where the model overemphasizes specific relationships found in the training data and becomes less generalizable to new areas. Hyperparameter tuning was conducted on the inner training split to provide for model optimization. This involved adjusting model tuning hyperparameters to converge model error metrics of the inner training split, and generally resulted in more accurate predictions of the outer novel testing data. In the model of BGB, only predictors with a feature importance score above 0.005 were retained.

S1.2.1. Remote Sensing Data

BERM relies on Landsat-8 and -9 Collection 2, Level 2, Tier 1 surface reflectance products as remote sensing data, which are freely available from the US Geological Survey and accessible through the Google Earth Engine web platform (Gorelick et al. 2017). Sensors aboard Landsat-8 and -9 are alike, allowing us to merge products seamlessly (Masek et al. 2020). Landsat data corresponding to field sampling dates were downloaded, screened for clouds with the Landsat 'QA_PIXEL' mask (Foga et al. 2017), screened for flooding with an in-house flooding model (O'Connell et al. 2021), and quality checked. Landsat-8 images are available dating back to 2013, and Landsat-9 images back to late 2021. These Landsat mission data are on a 30 x 30m spatial grid with at least 16 day returns; the two satellites are in offsetting orbits, effectively reducing the joined return period to 8 days from late 2021 onwards.

Remote sensing indices can help describe vegetation through spectral reflectance patterns (Rouse et al. 1974, Gitelson et al. 2002, Mishra and Ghosh 2015). We calculated a number of reflectance indices to use as model predictors alongside Landsat-8 and -9 bands 1 through 7 later in the modeling framework for aboveground vegetation models, one of which, NDVI, was also used in the BGB model (Table S3). Bands 1 through 7 cover the visible, near-infrared (NIR), and shortwave infrared portions of the electromagnetic spectrum (Masek et al. 2020). The indices we calculated and applied utilize a wider range of information across the electromagnetic spectrum and thus can be more informative than individual bands. For example, NDVI applies reflectance in NIR and red bands to assess vegetation biomass and health (Rouse et al. 1974, Running 1990).

#### S1.2.2. Advancing BERM to Version 2.0

To advance BERM from version 1.0 to version 2.0, the calibration dataset was expanded. This larger ground-truth field dataset was used to update and regenerate the above- and belowground models described in O'Connell et al. (2021). We rebuilt the models for each of the four aboveground plant biophysical variables measured: AGB, CHL, foliar N, and LAI, based on Landsat-8 and -9 surface reflectance and associated vegetation indices as predictors. Once aboveground XGB models were created, predictions were made for each aboveground variable on all dates in the interpolated field dataset. In addition to remote sensing data, the belowground vegetation model utilized aboveground vegetation predictions (from the aboveground models), climate data, and tide data as predictors. Data were joined on a monthly timestep. Time-series lags, rolling means, and differences of these variables were also calculated for potential inclusion as predictors.

To deduce the impact of the larger and more diverse calibration dataset, performance metrics were generated for the v1.0 models with the updated, debugged, and optimized model framework (as opposed to comparing to those reported in O'Connell et al. 2021), but with a limited calibration dataset, akin to that of O'Connell et al. (2021). These were then compared to the performance metrics generated for the v2.0 models, with the full calibration dataset.

#### *S1.3. Equations for model performance metrics*

Model performance metrics were calculated against the testing data with the following equations:

Equation S2.1.: $MBE = \sum y_i - \hat{y}_i / N$

Equation S2.2.: $RMSE = \sqrt{\sum (y_i - \hat{y}_i)^2 / N}$

Equation S2.3.: $nRMSE = \sqrt{\sum (y_i - \hat{y}_i)^2 / N} / (y_{max} - y_{min})$

Equation S2.4.: $COVRMSE = \sqrt{\sum (y_i - \hat{y}_i)^2 / N} / y_{mean}$

Pearson's correlation coefficient (*r*) was calculated using the 'stats' package in R.

*S1.4. BERMv2.0 Demonstration*

We illustrated BERM capabilities through a model application demonstration at the GCE-LTER Flux Tower Marsh on Sapelo Island, GA. This is a *S. alterniflora*-dominated marsh with extensive research activities through the GCE-LTER, and encompassed two of our nine sampling sites. Predictor data for model application included that used to build BERM (Landsat-8 and -9, Daymet, the Fort Pulaski NOAA tide gauge), as well as the USGS 3DEP 1 m digital elevation model (DEM) (U.S. Geological Survey 2023), which we resampled to 2 m. We spatially joined all data to 30 m Landsat pixel footprints for each model application date, and selected *S. alterniflora* locations for application by subsetting to 30 m pixel footprints which contained at least 95% cover of *S. alterniflora* via a species cover map developed by Alexander and Hladik (2015). Landscape estimates of BGB are outside of the scope of this study and are forthcoming in a future effort.

### Section S2: Results

#### *S2.1. Expanding the BERM calibration dataset*

The expansion of the calibration dataset resulted in increased sample sizes and altered shapes of the distributions of the calibration datasets for each of the BERM variables (Figure S4, Tables S3, S4). Among the five plant productivity variables, the size of the v2.0 calibration datasets were on average 466% larger than the v1.0s. The v2.0 calibration dataset was generated from a total of 6633 measurements, an increase of 3886 (141%) over the v1.0 calibration dataset (Table S2). The focus on additional sites, along with a longer period of interpolation between data collection (typically three months for newly collected calibration data, versus one to two month intervals in O'Connell et al. (2021)), accounts for the discrepancy between the increase of the data collection and the calibration dataset sizes. The range of the measurements used to build the v2.0 calibration dataset was expanded for each variable, and was on average 40% larger than that of the v1.0 calibration dataset (ABG: 40%, CHL: 77%, foliar N: 58%, LAI: 63%, BGB: 2%).

The mean AGB, CHL, foliar N, and LAI were greater, while the mean BGB was less in the v2.0 datasets (Wilcoxon rank sum tests, $p < 0.001$). The calibration dataset of AGB retained a similar shape after BERMv2.0 expansion, while those of LAI and BGB became more left skewed, and those of CHL and foliar N shifted from a right skew to a left skew. All v2.0 datasets became more heavy-tailed compared to the v1.0s.

Characteristics of the additional sites varied widely. At the four original sites, sample plot elevation within sites averaged 0.08m (standard deviation: 0.04 m), while sample plot elevation of the five additional sites was 0.38m (standard deviation: 0.14m). The range and standard deviation of the average root:shoot ratio were also both larger compared to original BERM sites (range: 2.53 and 3.94, standard deviation: 1.14 and 1.88, for original and additional BERM data collection sites, each respectively). Additional sites accounted for the lowest average BGB, LAI, and flooding frequency, highest average CHL and foliar N, and highest and lowest average AGB (Table S4). They also filled a wide gap in average BGB, where the four original sites had either below 516 $gm^{-2}$ or above 1474 g $m^{-2}$ BGB. Of the five additional sites, four had an average BGB within this range.

*S2.2. Improvement of BERMv2.0 Performance*

Performance metrics against the testing data of BERMv2.0 showed improvements in performance of the v2.0 model in some metrics (nRMSE), but other metrics showed a varied response (MBE, RMSE, COV RMSE, Correlation; Table S5). The MBE of v2.0 compared to v1.0 models was slightly smaller in magnitude for foliar N, CHL, and BGB, and larger in magnitude for AGB and LAI. The RMSE of AGB was larger, of foliar N, LAI, and BGB was smaller, and of CHL was similar in v2.0 models compared to v1.0. But, due to a larger range of calibration data in v2.0, the nRMSE was smaller, indicating more accurate predictions after the datasets were normalized in all v2.0 models compared to their v1.0 counterparts. In the model of BGB, the RMSE and nRMSE decreased by 15% and 28%, respectively, from the v1.0 to v2.0 models. The COV RMSE was consistently smaller in the v2.0 models with the exception of BGB. Correlation among field measurements and model predictions in the testing data was larger in v2.0 for LAI, CHL, and BGB, but smaller for AGB and foliar N.

*S2.3. Identifying over- and underprediction thresholds*

To identify thresholds of BERM BGB under- and overprediction, we created a linear model of prediction residual (observation - prediction) against observation. The observed BGB where the absolute value of the predicted residual through the linear model was identified as a threshold of under- or overprediction. No overprediction threshold was identified. An underprediction threshold was identified at an observed BGB of 1981 g $m^{-2}$ (Figure S6).

**Tables**

Table S1. Months and years of plot level data collection by site and variable, with counts indicated in parentheses.

| Site and Year | AGB | Foliar CHL | Foliar N | LAI | BGB |
|---|---|---|---|---|---|
| Dean Creek (deanc) | | | | | |
| 2021 | Jun, Aug, Nov (9) | Jun, Aug, Nov (9) | Jun, Aug, Nov (9) | Jun, Aug, Nov (9) | Jun, Aug, Nov (9) |
| 2022 | Feb, May, Aug, Nov (9) | Feb, May, Aug, Nov (9) | Feb, May, Aug, Nov (9) | Feb, May, Aug, Nov (9) | Feb, May, Aug, Nov (9) |
| 2023 | Feb (9) | Feb (9) | Feb (9) | Feb (9) | Feb (9) |
| Duplin River (dupli) | | | | | |
| 2021 | Jun, Aug, Nov (9) | Jun, Aug, Nov (9) | Jun, Aug, Nov (9) | Jun, Aug, Nov (9) | Jun, Aug, Nov (9) |
| 2022 | Feb, May, Aug, Nov (9) | Feb, May, Aug, Nov (9) | Feb, May, Aug, Nov (9) | Feb, May, Aug, Nov (9) | Feb, May, Aug, Nov (9) |
| 2023 | Feb (9) | Feb (9) | Feb (9) | Feb (9) | Feb (9) |
| Flux Tower Marsh A (fluxa) | | | | | |
| 2013 | Jun-Dec (18) | | | | |
| 2014 | Jan-Dec (18) | | Apr (6); May, Jul (18); Jun, | | |

| Site and Year | AGB | Foliar CHL | Foliar N | LAI | BGB |
| --- | --- | --- | --- | --- | --- |
| | | | Aug, Sep (17), Oct (20) | | |
| 2015 | Jan-Dec (18) | | Apr, Jul, Sep (18); Aug (17) | | |
| 2016 | Jan-Dec (18) | May, Jul-Oct (18) | May, Jul-Oct (9) | May, Jul, Aug (18); Oct (1) | May (5); Jul-Oct (9) |
| 2017 | Jan-May (18); Jun-Oct, Dec (20); Nov (40) | | | | May-Aug, Dec (6); Nov (12) |
| 2018 | Jan-Dec (20) | | | Aug, Nov (20) | Jan-Dec (6) |
| 2019 | Jan, Jul, Nov (40); Feb-May, Aug, Sep, Dec (20) | | | May, Aug, Dec (20) | Jan, Nov (12); Feb-Sep, Dec (6) |
| 2020 | Jan, Apr-Dec (20); Mar (40) | | | Sep (20) | Jan, Apr-Dec (6); Mar (12) |
| 2021 | Jan-Oct, Dec (20) | | | Jun, Jul (18) | Jan-Oct, Dec (6) |
| 2022 | Jan (40); Feb-Dec (20) | | | | Mar, Jun, Sep, Dec (6) |
| 2023 | Feb, Mar (20) | | | | Mar (6) |
| Flux Tower Marsh B (fluxb) | | | | | |
| 2016 | May, Jul-Oct (9) | May, Jul-Oct (9) | May, Jul-Oct (9) | May, Jul, Aug, Oct (9) | May, Jul-Oct (9) |
| 2021 | Jun, Aug, Nov (9) | Jun, Aug, Nov (9) | Jun, Aug, Nov (9) | Jun, Aug, Nov (9) | Jun, Aug, Nov (9) |

| Site and Year | AGB | Foliar CHL | Foliar N | LAI | BGB |
|---|---|---|---|---|---|
| 2022 | Feb, May, Aug, Nov (9) | Feb, May, Aug, Nov (9) | Feb, May, Aug, Nov (9) | Feb, May, Aug, Nov (9) | Feb, May, Aug, Nov (9) |
| 2023 | Feb (9) | Feb (9) | Feb (9) | Feb (9) | Feb (9) |
| Folly River (folly) | | | | | |
| 2021 | Jun, Aug, Nov (9) | Jun, Aug, Nov (9) | Jun, Aug, Nov (9) | Jun, Aug, Nov (9) | Jun, Aug, Nov (9) |
| 2022 | Feb, May, Aug, Nov (9) | Feb, May, Aug, Nov (9) | Feb, May, Aug, Nov (9) | Feb, May, Aug, Nov (9) | Feb, May, Aug, Nov (9) |
| 2023 | Feb (9) | Feb (9) | Feb (9) | Feb (9) | Feb (9) |
| Hunt Camp (huntc) | | | | | |
| 2021 | Jun, Aug, Nov (9) | Jun, Aug, Nov (9) | Jun, Aug, Nov (9) | Jun, Aug, Nov (9) | Jun, Aug, Nov (9) |
| 2022 | Feb, May, Aug, Nov (9) | Feb, May, Aug, Nov (9) | Feb, May, Aug, Nov (9) | Feb, May, Aug, Nov (9) | Feb, May, Aug, Nov (9) |
| 2023 | Feb (9) | Feb (9) | Feb (9) | Feb (9) | Feb (9) |
| North Sapelo (north) | | | | | |
| 2021 | Jun, Aug, Nov (9) | Jun, Aug, Nov (9) | Jun, Aug, Nov (9) | Jun, Aug, Nov (9) | Jun, Aug, Nov (9) |
| 2022 | Feb, May, Aug, Nov (9) | Feb, May, Aug, Nov (9) | Feb, May, Aug, Nov (9) | Feb, May, Aug, Nov (9) | Feb, May, Aug, Nov (9) |
| 2023 | Feb (9) | Feb (9) | Feb (9) | Feb (9) | Feb (9) |
| Skidaway (skida) | | | | | |
| 2016 | May, Jul-Oct (9) | May, Jul-Oct (9) | May, Jul-Oct (9) | May, Jul, Aug, Oct (9) | May, Jul-Oct (9) |
| 2021 | Jun, Aug, | Jun, Aug, | Jun, Aug, | Jun, Aug, | Jun, Aug, |

| Site and Year | AGB | Foliar CHL | Foliar N | LAI | BGB |
|---|---|---|---|---|---|
| | Nov (9) | Nov (9) | Nov (9) | Nov (9) | Nov (9) |
| 2022 | Feb, May, Aug, Nov (9) | Feb, May, Aug, Nov (9) | Feb, May, Aug, Nov (9) | Feb, May, Aug, Nov (9) | Feb, May, Aug, Nov (9) |
| 2023 | Feb (9) | Feb (9) | Feb (9) | Feb (9) | Feb (9) |
| UGAMI (ugami) | | | | | |
| 2016 | May, Jul-Oct (9) | May, Jul-Oct (9) | May, Jul-Oct (9) | May, Jul, Aug, Oct (9) | May, Jul-Oct (9) |
| 2021 | Jun, Aug, Nov (9) | Jun, Aug, Nov (9) | Jun, Aug, Nov (9) | Jun, Aug, Nov (9) | Jun, Aug, Nov (9) |
| 2022 | Feb, May, Aug, Nov (9) | Feb, May, Aug, Nov (9) | Feb, May, Aug, Nov (9) | Feb, May, Aug, Nov (9) | Feb, May, Aug, Nov (9) |
| 2023 | Feb (9) | Feb (9) | Feb (9) | Feb (9) | Feb (9) |

[a]Aboveground Biomass; [b]Chlorophyll; [c]Nitrogen; [d]Leaf Area Index; [e]Belowground Biomass;

Table S2. Plot-level measurement counts by model version.

| Model | AGB[a] | Foliar CHL[b] | Foliar N[c] | LAI[d] | BGB[e] |
|---|---|---|---|---|---|
| BERMv1.0 | 1,581 | 225 | 328 | 263 | 350 |
| BERMv2.0 | 2,937 | 801 | 901 | 895 | 1,099 |

[a]Aboveground Biomass; [b]Chlorophyll; [c]Nitrogen; [d]Leaf Area Index; [e]Belowground Biomass;

Table S3. Remote sensing indices applied as BERMv2.0 predictor variables.

| Index | Equation | Reference |
|---|---|---|
| GARI | $\left(B5-\left(B3-1.7*\left(B2-B4\right)\right)\right)/\left(B5+\left(B3-1.7*\left(B2-B4\right)\right)\right)$ | Gitelson et al. (1996) |

| Index | Equation | Reference |
|---|---|---|
| GCI | $(B5/B3)-1$ | Gitelson et al. (2003) |
| GNDVI | $(B5-B3)/(B5+B3)$ | Gitelson et al. (1996) |
| MTVI1 | $1.2*(1.2*(B5-B3)-2.5*(B4-B3))$ | Haboudane et al. (2004) |
| MTVI2 | $(1.5*(1.2*(B5-B3)-2.5*(B4-B3)))/((2*B5+1)^2-6*B5-5*B4^{0.5})-0.5^{0.5}$ | |
| NDMI | $(B1-B6)/(B1+B6)$ | Hardisky et al. (1983) |
| NDVI | $(B5-B4)/(B5+B4)$ | Rouse et al. (1974) |
| Pheno | $(B5-B6)/(B5+B6)$ | O'Connell et al. (2017) |
| RDVI | $(B5-B4)/(B5+B4)^{0.5}$ | Roujean and Breon (1995) |
| VARI | $(B3-B4)/(B2+B3+B4)$ | Gitelson et al. (2002) |

Table S4. Mean (and range) values of pixel level ground-truth data collection and scaling-up data which serve as predictors for our belowground biomass model.

| Variable | Dean Creek | Duplin River | Flux A | Flux B | Folly River | Hunt Camp | North Sapelo | Skidaway | UGA MI |
|---|---|---|---|---|---|---|---|---|---|
| BGB[1a] | 735 (329-1742) | 1358 (786-2038) | 1213 (119-3928) | 509 (117-1718) | 1075 (501-1680) | 472 (243-938) | 771 (167-1610) | 510 (206-1014) | 1520 (531-3036) |
| AGB[1b] | 417 (168-821) | 262 (150-412) | 293 (33-1120) | 153 (40-299) | 227 (122-395) | 285 (56-584) | 139 (68-221) | 276 (124-532) | 339 (149-508) |
| LAI[c] | 1.40 (0.86-2.99) | 1.00 (0.47-2.37) | 1.47 (0.36-5.76) | 0.45 (0.14-1.19) | 0.53 (0.21-1.19) | 0.67 (0.30-1.74) | 0.38 (0.16-0.65) | 0.75 (0.39-1.23) | 1.33 (0.73-3.25) |
| % foliar | 1.37 | 1.27 | 1.14 | 1.32 | 1.35 | 1.49 | 1.38 | 1.16 | 1.24 |

| Variable | Dean Creek | Duplin River | Flux A | Flux B | Folly River | Hunt Camp | North Sapelo | Skidaway | UGA MI |
|---|---|---|---|---|---|---|---|---|---|
| N[d] | (0.92-2.00) | (0.97-1.70) | (0.79-1.58) | (0.68-1.93) | (1.01-1.84) | (1.08-2.01) | (1.17-1.92) | (0.52-1.92) | (0.72-1.90) |
| Foliar CHL[2e] | 1.05 (0.71-1.43) | 0.93 (0.73-1.23) | 0.65 (0.44-0.87) | 0.87 (0.44-1.13) | 0.99 (0.82-1.19) | 1.11 (0.75-1.32) | 1.03 (0.77-1.34) | 0.99 (0.49-1.38) | 0.95 (0.60-1.20) |
| Green-up day of year[f] | 29 (11-53) | 43 (29-56) | 39 (27-53) | 39 (28-53) | 43 (29-55) | 37 (27-50) | 40 (28-53) | 41 (28-55) | 40 (28-53) |
| Elevation[3] | 0.80 (0.69-0.90) | 0.97 (0.91-1.07) | 0.74 (0.65-0.82) | 0.72 (0.69-0.75) | 0.89 (0.89-0.91) | 0.59 (0.54-0.63) | 0.83 (0.77-0.89) | 0.79 (0.72-0.87) | 0.80 (0.75-0.85) |
| Inundation intensity[g] | 0.05 (0.01-0.13) | 0.02 (-0.00-0.05) | 0.05 (0.02-0.10) | 0.14 (0.06-0.30) | 0.04 (0.01-0.09) | 0.10 (0.05-0.19) | 0.10 (0.02-0.22) | 0.13 (0.04-0.24) | 0.06 (0.01-0.12) |
| Dry intensity[h] | 1.39 (1.05-1.69) | 1.62 (1.34-1.93) | 1.39 (1.09-1.59) | 1.04 (0.71-1.33) | 1.44 (1.22-1.68) | 1.18 (0.91-1.41) | 1.18 (0.85-1.49) | 1.05 (0.80-1.32) | 1.37 (1.08-1.68) |
| Flooding frequency[i] | 0.14 (0.05-0.21) | 0.08 (0.03-0.11) | 0.11 (0.07-0.15) | 0.32 (0.22-0.46) | 0.15 (0.09-0.18) | 0.17 (0.13-0.24) | 0.28 (0.19-0.38) | 0.35 (0.28-0.42) | 0.15 (0.07-0.21) |
| Monthly mean max. temp.[3] | 26 (14-35) | 26 (14-35) | 26 (14-35) | 26 (14-35) | 26 (14-35) | 26 (14-35) | 26 (14-35) | 26 (14-35) | 26 (14-35) |
| Monthly mean min. temp.[3] | 16 (4-25) | 16 (3-25) | 16 (3-25) | 16 (3-25) | 16 (3-24) | 16 (3-25) | 16 (3-25) | 15 (2-24) | 16 (4-25) |
| Monthly mean precip.[2] | 3.62 (0.15-9.64) | 3.65 (0.16-9.71) | 3.69 (0.16-9.62) | 3.71 (0.16-9.57) | 3.70 (0.21-10.02) | 3.75 (0.15-9.63) | 3.82 (0.13-10.01) | 3.71 (0.18-11.46) | 3.64 (0.16-9.67) |

[1]g m$^{-2}$; [2]mg g$^{-1}$; [3]m NAVD88; [3]° C; [2]mm day$^{-1}$; [a]Belowground Biomass; [b]Aboveground Biomass; [c]Leaf Area Index; [d]Nitrogen; [e]Chlorophyll; [f]Day of Year; [g]Inundation Intensity; [h]Dry Intensity; [i]Flooding Frequency;

Table S5. Testing data mean goodness of fit metrics, including Mean Bias Error (MBE; units of the variable), Root Mean Squared Error (RMSE; units of the variable), normalized RMSE (nRMSE), coefficient of variation of RMSE (COV RMSE), and Correlation (*r*) for the *Spartina alterniflora* aboveground vegetation Extreme Gradient Boosting models. Sample size for training and testing data also are provided (N-train and N-test, respectively). Variable abbreviations: AGB (Aboveground Biomass), CHL (Foliar Chlorophyll), Foliar N (Foliar Nitrogen), and LAI (Leaf Area Index).

| Model | Variable[1] | MBE[1a] | RMSE[b] | nRMSE[c] | COV RMSE[d] | Correlation | *n*-Test | *n*-Train |
|---|---|---|---|---|---|---|---|---|
| BERMv1.0 | AGB[1] | 0.15 | 104.09 | 14.7% | 44.7% | 0.68 | 231 | 433 |
| BERMv2.0 | AGB[1] | 1.07 | 110.01 | 11.6% | 42.4% | 0.63 | 832 | 1,417 |
| BERMv1.0 | Foliar N[2] | -0.14 | 0.25 | 20.9% | 23.0% | 0.54 | 79 | 123 |
| BERMv2.0 | Foliar N[2] | -0.01 | 0.24 | 16.4% | 17.9% | 0.44 | 509 | 781 |
| BERMv1.0 | LAI[3] | 0.00 | 0.47 | 21.1% | 50.7% | 0.47 | 62 | 140 |
| BERMv2.0 | LAI[3] | 0.06 | 0.36 | 9.6% | 44.8% | 0.65 | 499 | 819 |
| BERMv1.0 | CHL[4] | -0.09 | 0.15 | 33.8% | 21.5% | 0.27 | 59 | 106 |
| BERMv2.0 | CHL[4] | 0.00 | 0.15 | 15.2% | 14.8% | 0.58 | 490 | 763 |

[1]$g\ m^{-2}$; [2]%; [3]$m\ m^{-1}$; [4]$mg\ g^{-1}$; [1]$g\ m^{-2}$; [a]Mean Bias Error; [b]Root Mean Square Error; [c]Normalized Root Mean Square Error; [d]Coefficient of Variation of Root Mean Square Error;

Table S6. Regression Commonality Analysis results, with commonality coefficients that depict explained variation in a multiple regression analysis of each variable by Site, Year, and the interaction between Site and Year. '% Total' refers to the percent of total explained variation associated with that grouping.

| Belowground Biomass | Commonality Coefficient | | | % Total | | |
|---|---|---|---|---|---|---|
| | Site | Year | Site:Year | Site | Year | Site:Year |
| Observed | 0.28 | 0.02 | 0.03 | 84 | 6 | 10 |
| Predicted | 0.29 | 0.02 | 0.03 | 85 | 5 | 10 |

**Figures**

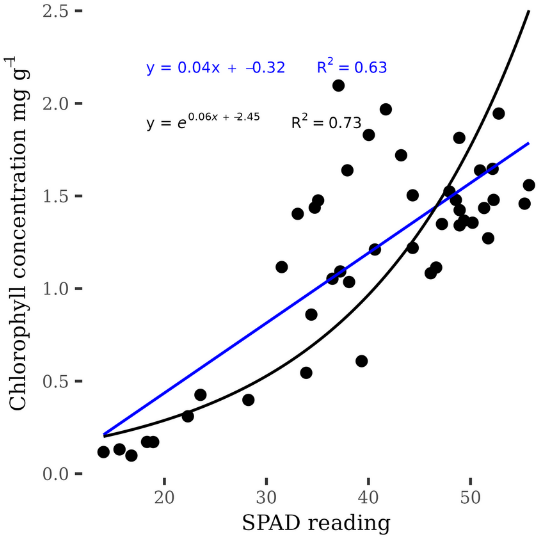


*Figure S1. SPAD measurements and leaf chlorophyll concentrations, with linear (black line and equation) and exponential (blue line and equation) prediction models.*

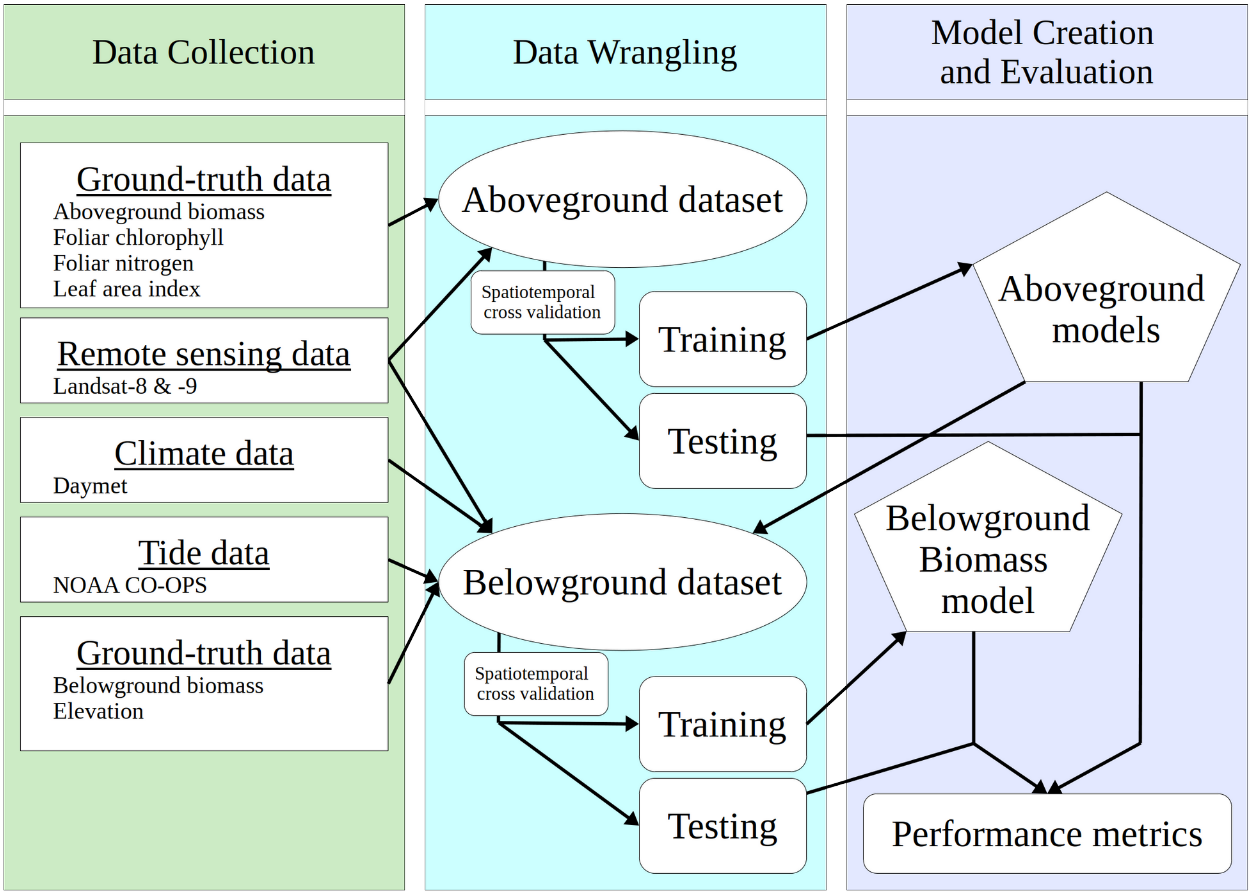


*Figure S2. BERM workflow, including data sources for candidate predictors and modeling organization.*

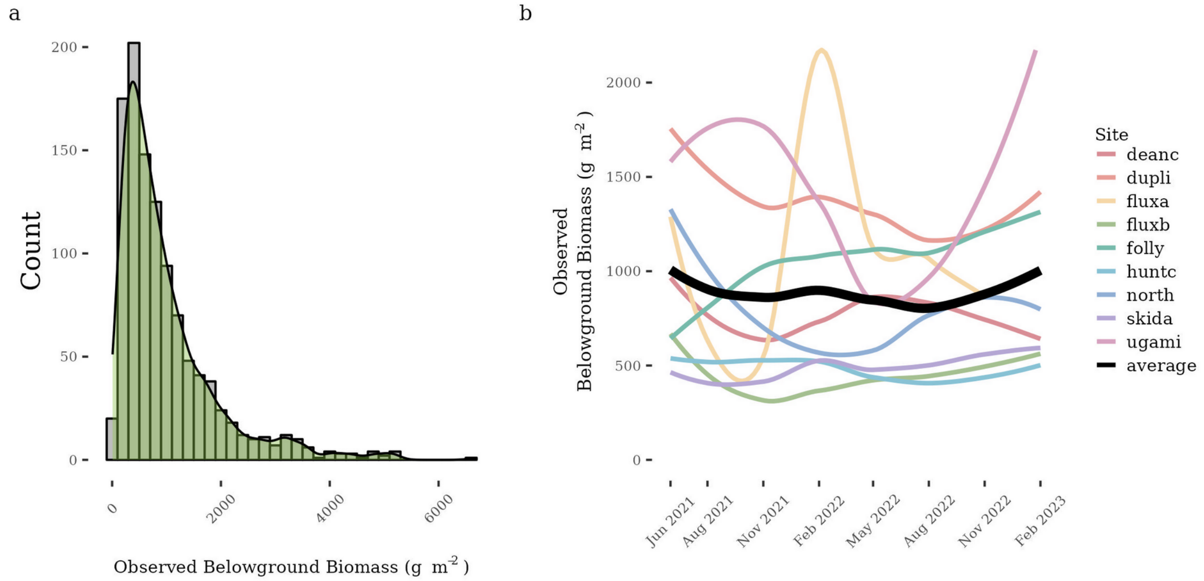


*Figure S3. a). Plot-level observed BGB. b) Smoothed site level observed belowground biomass by sample date from 2021 to 2023.*

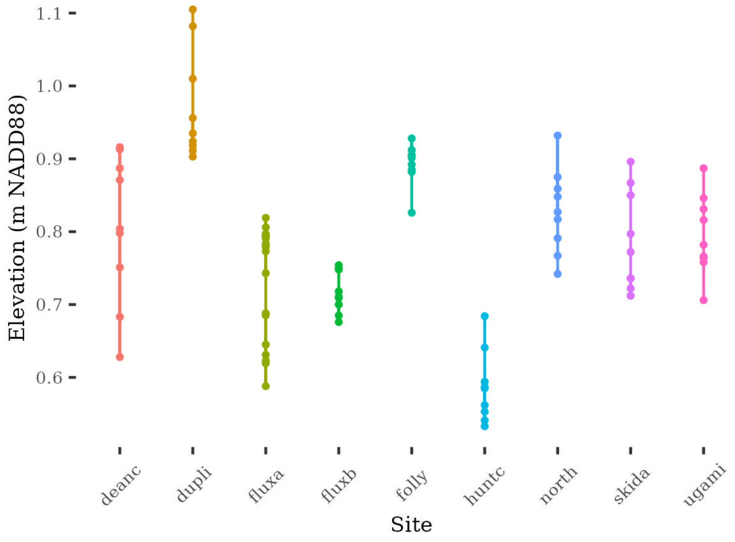


*Figure S4. Plot-level elevation by site.*

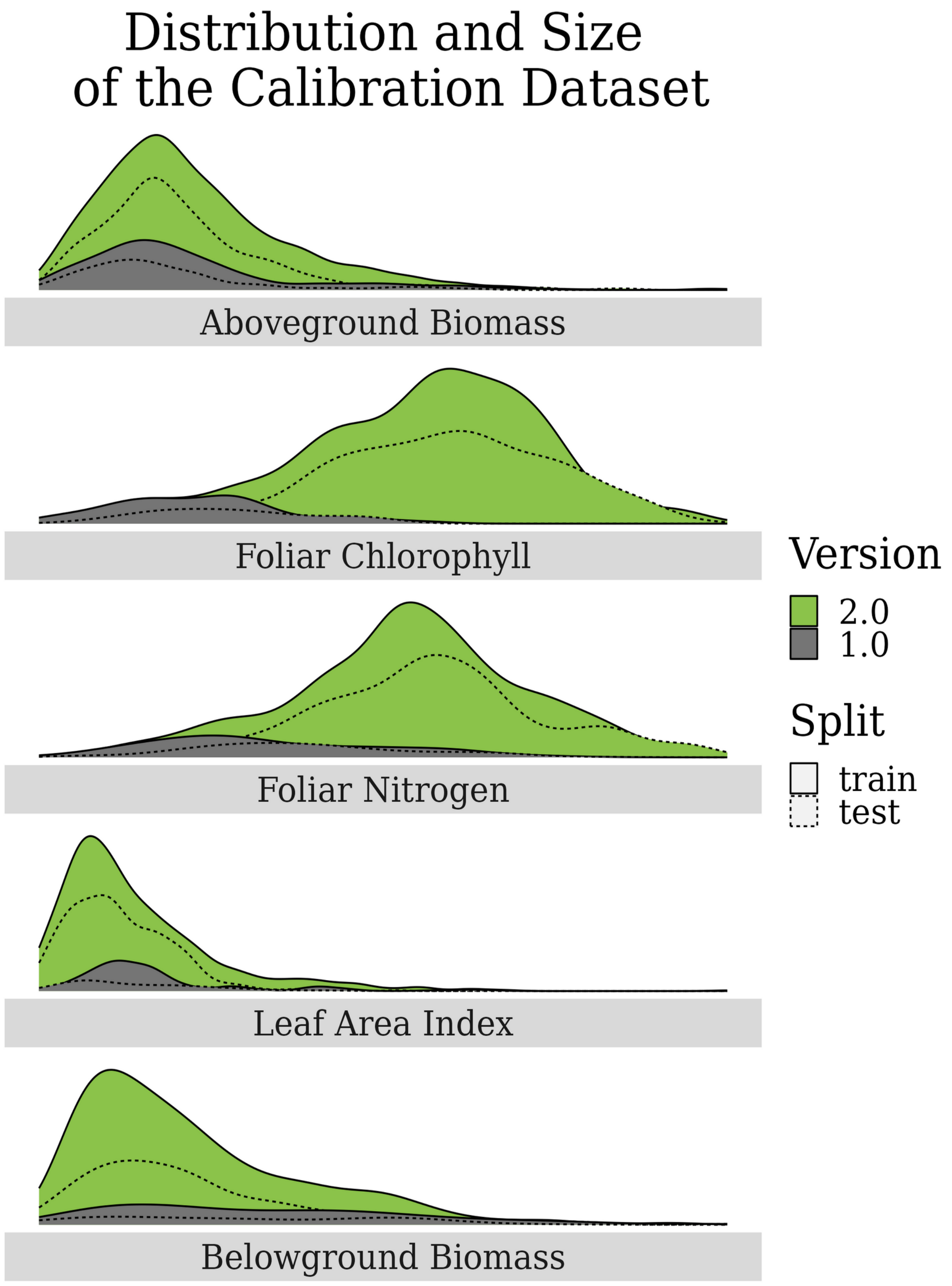


*Figure S5. Distribution and relative size of the v1.0 (gray) and v2.0 (green) calibration dataset (pixel-level). Model-building training and testing splits*

*are depicted by solid and dotted lines, respectively. For each variable, the calibration dataset was expanded in size and for some, expanded in range.*

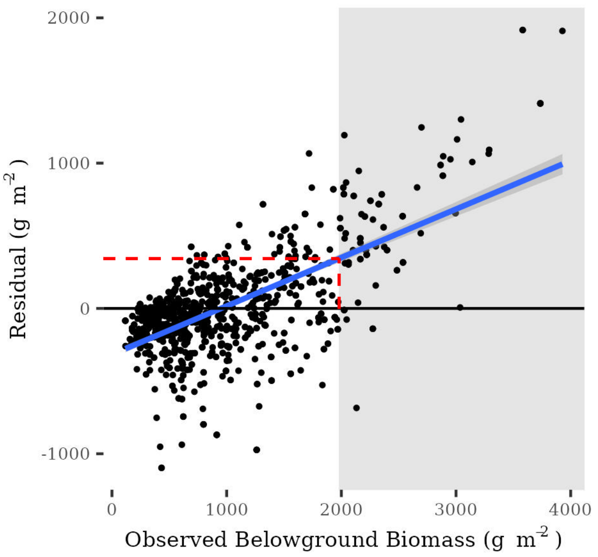


*Figure S6. Linear model of BGB prediction residuals against observations. The blue line depicts the linear model. The horizontal red dashed line depicts the RMSE (344 g m$^{-2}$), and the vertical red line refers to the observed belowground biomass where the linear model reaches this as residual.*
